# Mammalian musculoskeletal regeneration is associated with reduced inflammatory cytokines and an influx of T cells

**DOI:** 10.1101/723783

**Authors:** Thomas R. Gawriluk, Jennifer Simkin, Corin K. Hacker, John M. Kimani, Stephen G. Kiama, Vanessa O. Ezenwa, Ashley W. Seifert

## Abstract

Whether the immune response to injury contributes to tissue regeneration is not well understood. We quantified systemic and local cytokines during ear pinna repair to provide the first comprehensive comparison of the immune response to injury between mammalian regeneration (*A. cahirinus* and *A. percivali*) and fibrotic repair (*M. musculus*). Importantly, by comparing laboratory-reared and wild-caught animals we identified responses specifically associated with healing outcome. Fibrotic repair showed a greater local release of IL-6, CCL2 and CXCL1. Conversely, regeneration showed decreased circulating IL-5, IL-6, IL-17, CCL3 and CXCL1 and increased local IL-12 and IL-17. The differential IL-6 response was substantiated by increased pSTAT3 during the inflammatory phase of fibrotic repair and with blastema formation and tissue morphogenesis in *Acomys*. COX-2 inhibition was not sufficient to induce regeneration. Interestingly, a unique influx of lymphocytes was coupled with regeneration and RNA-expression analysis suggested they were regulatory T cells. Together, the data support regeneration-specific inflammation and T cell responses in *Acomys*.

## INTRODUCTION

Epimorphic regeneration in response to tissue damage occurs as a chronological and overlapping series of processes that includes hemostasis, inflammation, re-epithelialization, activation of local progenitor cells, tissue morphogenesis, and replacement of the injured tissue. The outcome of these processes is scar-free healing. In contrast, most human injuries heal by fibrotic repair, which is characterized by limited cellular proliferation and intense collagen deposition that results in a scar repairing the damaged tissue [1]. As with any trauma or infection that disrupts tissue architecture, regeneration and fibrotic repair are concomitant with a multiphasic immune response that promotes hemostasis, creates inflammation, protects against microbial infection, initiates re-epithelialization, and stimulates a local fibrotic response [2]. During most instances of regeneration (e.g., limb, fin, digit tip, etc.) there is an apparent resolution of acute inflammation that coincides with the accumulation of resident cells, which subsequently re-enter the cell cycle and self-organize to undergo morphogenesis [3, 4]. This transition from an inflammatory environment to morphogenesis is synonymous with regenerative blastema formation [5]. As such, the injured tissue must precisely coordinate dynamic interactions between cells and factors (i.e., cytokines, chemokines, growth factors etc.) within the injury microenvironment to resolve the inflammatory response and promote blastema formation.

Despite a rich literature describing the effects of immune cells and their products in non-regenerating wounds [6, 7], our knowledge of the immune response during epimorphic regeneration remains poor [8–10]. Recent studies in fish, frogs, salamanders, and spiny mice support that immune cells and their products are required for blastema formation and successful regeneration. For instance, complete or timed depletion of macrophages or regulatory T cells (T_REG_) prevents re-epithelialization and subsequent blastema formation [11–18] and blocking reactive oxygen species (ROS) production elicits a similar outcome [19–21]. Perhaps not unexpectedly, similar experiments in non-regenerating systems cause incomplete wound closure and angiogenesis, suggesting that the same immune signals initiate fibrotic repair and regeneration [22–27]. Moreover, comparing the immune response to injury between fetal and adult mammals [28–30], pre- and post-metamorphic amphibians [31–33], closely-related regenerating and non-regenerating vertebrates [34], and regeneration-competent and scarring tissues in the same animal [35–37] all support that reduced inflammation and a muted immune response promotes epimorphic regeneration over fibrotic repair. However, an important series of studies support the idea that some immune cells are passive participants during tissue regeneration. For example, removal of the spleen [38], or induction of leukopenia [39] during newt limb regeneration demonstrate that a severely reduced leukocyte response does not prevent blastema formation or regeneration.

These contrasting viewpoints raise several unanswered questions. (1) Are there specific factors produced by immune cells that polarize local cell phenotypes and specifically promote regeneration or fibrotic repair? (2) Does the inflammatory response impede blastema formation and subsequent regeneration in adult mammals? (3) Are the initial stages of fibrotic repair and regeneration driven by different immune responses, such that altering the immune response could stimulate regeneration in lieu of fibrotic repair? In order to address these questions and to test how immune cell factors specifically affect resident cells in injured tissue, there is a need to establish the timing and extent to which these factors are deployed during regeneration and fibrotic repair.

In this study, we provide insight of the immune mechanisms that coincide with mammalian regeneration by following-up on the relatively recent discovery that multiple species of spiny mice (e.g., *Acomys cahirinus, A. percivali, A. kempi*) regenerate skin and musculoskeletal tissue [15, 40–42]. Previous work in spiny mice supports that epimorphic regeneration is associated with resident cell activation, cell cycle progression, cell proliferation, and an inflammatory response distinguished by a prolonged signature of NADPH-oxidase-derived ROS [15, 40, 42]. Additionally, macrophages are required for blastema formation in spiny mice [15], and there is additional evidence that macrophage subtypes differentially drive regeneration or fibrotic repair in rodents [15–17]. Therefore, we sought to characterize and compare the cytokine response to injury during fibrotic repair and regeneration to specifically test if the immediate immune response to injury is different between these two healing outcomes.

We compared the injury response using a 4 mm ear punch assay among three species (*A. cahirinus, A. percivali and M. musculus*) and two source populations (wild-caught *A. percivali* and *M. musculus*, and laboratory-reared *A. cahirinus* and *M. musculus*) using a panel of sixteen cytokines. Our results showed that injury across all groups induced a common set of pro-inflammatory cytokines (IL-1ß, IL-6, TNFα) and leukocyte chemotactic factors (GM-CSF/CSF2, and MIP-1α/CCL3) with a similar timed resolution, supporting that some signals of acute inflammation are a shared feature of regeneration and fibrotic repair. Supporting a difference between regeneration and fibrotic repair, we observed differential responses for several pro-inflammatory cytokines during the acute inflammatory phase. Surprisingly, we found a faster, stronger and prolonged adaptive immune response characterized by T cell influx during regeneration.

## RESULTS

### Cross-species validation of cytokine detection in rodent serum and tissue

To begin characterizing the mammalian immune response during epimorphic regeneration, we analyzed sixteen cytokines (Interleukin 1-alpha (IL-1α), IL-1β, IL-2, IL-4, IL-5, IL-6, IL-10, IL-12p70, IL-17, chemokine (C-C motif) ligand 2 (CCL2) (a.k.a. monocyte chemoattractant protein 1 or MCP-1), CCL3 (a.k.a. macrophage inflammatory protein 1α or MIP-1α), CCL5 (a.k.a. regulated on activation, normal T cell expressed and secreted or RANTES), colony-stimulating factor 2 (CSF2) (a.k.a. granulocyte-macrophage colony-stimulatory factor or GM-CSF), tumor necrosis factor-alpha (TNFα), interferon-gamma (IFNγ) and chemokine (C-X-C motif) ligand 1 (CXCL1) (a.k.a. KC)) using a custom-designed sandwich ELISA array. We used this assay to compare five groups: three at the University of Kentucky, (1) laboratory-reared, outbred *Mus musculus* (Mm-UKY), (2) wild-caught *M. musculus* (Mm-Wild), (3) laboratory-reared *Acomys cahirinus* (Ac), and two at the University of Nairobi, (4) outbred *M. musculus* reared by a local breeder (Mm-Kenya) and (5) wild-caught *A. percivali* (Ap). Our experimental design allowed us to compare cytokine responses between regenerating and non-regenerating species (Ac and Ap compared to Mm-UKY, Mm-Kenya and Mm-Wild), and between immune-challenged and laboratory-reared animals (Mm-Kenya, Mm-Wild and Ap compared to Mm-UKY and Ac).

Parallelism analysis showed comparable slopes between *Mus* and *Acomys* serum and tissue samples with the mouse standard curve for a majority of cytokines (Table 1 and Supplementary Figure 1). Given non-parallel slopes, we did not validate using this assay to compare IL-10 or CCL5 between species (Figure S1). Several cytokines not present in *Acomys* serum were quantified in tissue lysate (Ac: IL-1β, IL-4, IL-6, IL-17, CSF2, CCL2; Ap: IL-17, CSF2, CXCL1) supporting that the serum concentration was below the limit of detection and that the antibody binding epitopes were conserved between species. Therefore, if the cytokine was detected in one tissue source or one *Acomys* species, we concluded it could be detected in the other source or species. This provided us with a way to determine if the cytokine was present or absent. A comparison of full-length predicted amino acid sequences between *A. cahirinus* and *M. musculus* indicated conservation—minimum of 56.8% (IFNγ) to a maximum of 95.8% (TNFα) (Table 2 and Supplemental Figure 2A-Q). Given that CXCL1 was the only cytokine not detected in the Ac samples, this supports that it was likely not being present versus a failure to detect it. Together, these results supported that the ELISA could be used to directly compare changes in most of the cytokines between species.

**Table 1:** Comparison of cytokine slopes from parallelism test of cytokine assay. This was used to determine which species and source comparisons could be made for each cytokine in the Comparison column.

| Antigen | Standard | Sample | *Mus musculus (Mm)* | *Acomys cahirinus (Ac)* | *Acomys percivali (Ap)* | Comparison |
| --- | --- | --- | --- | --- | --- | --- |
| IL-1α | -0.89 ± 0.03 | Serum:<br>Tissue: | -1.05 ± 0.03*<br>*Too few points*<sup>^</sup> | -0.82 ± 0.14*<br>-1.03 ± 0.04* | -1.37 ± 0.10*<br>-0.97 ± 0.18* | Mm, Ac, Ap<br>Mm, Ac, Ap |
| IL-1β | -0.93 ± 0.10 | Serum:<br>Tissue: | *Too few points*<sup>&</sup><br>-0.94 ± 0.02* | *Not detected*<br>-0.86 ± 0.03* | -1.63 ± 0.21*<br>-0.65 ± 0.07* | Mm, Ap<br>Mm, Ac, Ap |
| IL-2 | -1.34 ± 0.05 | Serum:<br>Tissue: | *Too few points*<sup>&</sup><br>-1.21 ± 0.10* | *Not detected*<br>-0.59 ± 0.22* | -0.87 ± 0.00*<br>-0.57 ± 0.20* | Mm, Ap<br>Mm, Ac, Ap |
| IL-4 | -0.89 ± 0.06 | Serum:<br>Tissue: | -0.88 ± 0.18*<br>-0.69 ± 0.09* | *Too few points*<sup>&</sup><br>-0.68 ± 0.39* | -2.16 ± 0.32*<br>-0.75 ± 0.12* | Mm, Ap<br>Mm, Ac, Ap |
| IL-5 | -1.13 ± 0.01 | Serum:<br>Tissue: | -1.63 ± 0.20*<br>-0.73 ± 0.05* | -1.53 ± 0.96*<br>*Too few points*<sup>&</sup> | -2.52 ± 0.28*<br>-0.87 ± 0.03* | Mm, Ac, Ap<br>Mm, Ap |
| IL-6 | -1.07 ± 0.02 | Serum:<br>Tissue: | -0.74 ± 0.12*<br>-0.80 ± 0.14* | *Too few points*<sup>&</sup><br>-0.47 ± 0.06* | -1.81 ± 0.11*<br>-0.63 ± 0.11* | Mm, Ap<br>Mm, Ac, Ap |
| IL-10 | -1.56 ± 0.03 | Serum:<br>Tissue: | *Not detected*<br>*Not detected* | *Not detected*<br>0.52 ± 0.25 | *Not detected*<br>0.24 ± 0.52 | None<br>None |
| IL-12 | -1.10 ± 0.02 | Serum:<br>Tissue: | -0.68 ± 0.16*<br>-0.73 ± 0.17* | -1.55 ± 0.49*<br>-0.68 ± 0.16* | -1.26 ± 0.18*<br>-0.63 ± 0.15* | Mm, Ac, Ap<br>Mm, Ac, Ap |
| IL-17 | -1.03 ± 0.02 | Serum:<br>Tissue: | -0.72 ± 0.25*<br>-1.28 ± 0.04* | -0.16 ± 0.23<br>-0.85 ± 0.34* | -0.49 ± 0.16<br>-0.81 ± 0.03* | Mm<br>Mm, Ac, Ap |
| CSF2 | -1.45 ± 0.05 | Serum:<br>Tissue: | *Not detected*<br>-0.71 ± 0.11* | *Not detected*<br>-0.68 ± 0.25* | *Too few points*<sup>&</sup><br>-0.70 ± 0.10* | None<br>Mm, Ac, Ap |
| CCL2 | -1.17 ± 0.00 | Serum:<br>Tissue: | -1.23 ± 0.14*<br>-0.97 ± 0.02* | *Too few points*<sup>&</sup><br>-1.23 ± 0.14* | -1.56 ± 0.13*<br>-0.63 ± 0.10* | Mm, Ap<br>Mm, Ac, Ap |
| CCL3 | -0.98 ± 0.06 | Serum:<br>Tissue: | -2.05 ± 0.03*<br>-0.93 ± 0.01* | -1.25 ± 0.09*<br>-0.95 ± 0.01* | -1.01 ± 0.09*<br>-0.94 ± 0.06* | Mm, Ac, Ap<br>Mm, Ac, Ap |
| CCL5 | -0.90 ± 0.07 | Serum:<br>Tissue: | -0.78 ± 0.03*<br>-0.90 ± 0.16* | *Not detected*<br>0.34 ± 0.13 | 0.33 ± 0.32<br>0.07 ± 0.27 | Mm<br>Mm |
| TNFα | -1.10 ± 0.06 | Serum:<br>Tissue: | -1.36 ± 0.38*<br>-1.14 ± 0.10* | -2.15 ± 0.01*<br>-0.93 ± 0.01* | -1.09 ± 0.17*<br>-1.21 ± 0.03* | Mm, Ac, Ap<br>Mm, Ac, Ap |
| IFNγ | -0.79 ± 0.10 | Serum:<br>Tissue: | -0.60 ± 0.16*<br>-0.81 ± 0.31* | -1.48 ± 0.72*<br>-0.72 ± 0.05* | -1.21 ± 0.22*<br>-0.84 ± 0.05* | Mm, Ac, Ap<br>Mm, Ac, Ap |
| CXCL1 | -1.20 ± 0.02 | Serum:<br>Tissue: | -1.15 ± 0.27*<br>-1.11 ± 0.05* | *Not detected*<br>*Too few points*<sup>&</sup> | *Not detected*<br>-1.15 ± 0.29* | Mm<br>Mm, Ap |

Slopes in *italics* are not different from zero ( $P > 0.05$ for regression test)
\* denotes slope is similar to the standard and can be reliably quantified
& denotes values are below the assay's lower limit of detection
^ denotes values are above the assay's upper limit of quantification

**Table 2:** Comparison of *A. cahirinus* predicted peptide sequences used in cytokine analysis

| Gene ID | Peptide ID | % similar to<br>*M. musculus* | % similar to<br>*R. norvegicus* | % similar to<br>*H. sapiens* |
| --- | --- | --- | --- | --- |
| *Il1a* | IL-1α | 82.90 | 86.90 | 66.90 |
| *Il1b* | IL-1β | 87.71 | 87.71 | 72.35 |
| *Il2* | IL-2 | 70.41 | 84.61 | 71.59 |
| *Il4* | IL-4 | 66.45 | 77.21 | 56.32 |
| *Il5* | IL-5 | 90.15 | 84.84 | 68.93 |
| *Il6* | IL-6 | 75.23 | 78.97 | 48.13 |
| *Il10* | IL-10 | 85.05 | 87.93 | 74.13 |
| *Il12a* | IL-12 p35 | 70.42 | 80.54 | 55.25 |
| *Il12b* | IL-12 p40 | 73.58 | 74.10 | 63.33 |
| *Il17a* | IL-17 | 84.88 | 83.13 | 75.58 |
| *Cxcl1* | KC | 80.37 | 81.30 | 69.15 |
| *Ccl2* | MCP-1 | 73.50 | 70.86 | 43.70 |
| *Ccl3* | MIP-1α | 93.54 | 94.62 | 81.72 |
| *Ccl5* | RANTES | 88.04 | 84.78 | 85.86 |
| *Csf2* | GM-CSF | 65.24 | 73.75 | 61.70 |
| *IFNγ* | IFNγ | 56.77 | 57.05 | 41.56 |
| *Tnf* | TNFα | 95.81 | 96.65 | 84.93 |

### Fibrotic repair is associated with elevated amounts of circulating IL-5, IL-6 and CCL3

Using our cytokine assay, we first compared circulating serum cytokine concentrations from uninjured animals among groups (species and source population) to establish a systemic baseline for each group (Figure 1A). A total of 13 cytokines were compared as CSF2 was not present in the serum of any species. While many baseline concentrations were similar between groups, immune-challenged animals (i.e., wild) exhibited higher IL-4, IL-6, CCL2, and TNFα compared to laboratory-reared animals (Figure 1A). Interestingly, the Mm-Kenya animals were a transitional group between Mm-UKY and Mm-Wild for TNFα and IL-4 (Figure 1A). Heightened concentrations of IL-6, TNFα and IL-4 support previous pathogen exposure and a possibility of current infection [43, 44]. Thus, Mm-Kenya, Mm-Wild and Ap had a relatively activated immune system, while Mm-UKY and Ac possessed a more naïve immune system [45]. There were no consistent differences between regenerators and non-regenerators (Figure 1A).

**Figure 1.**
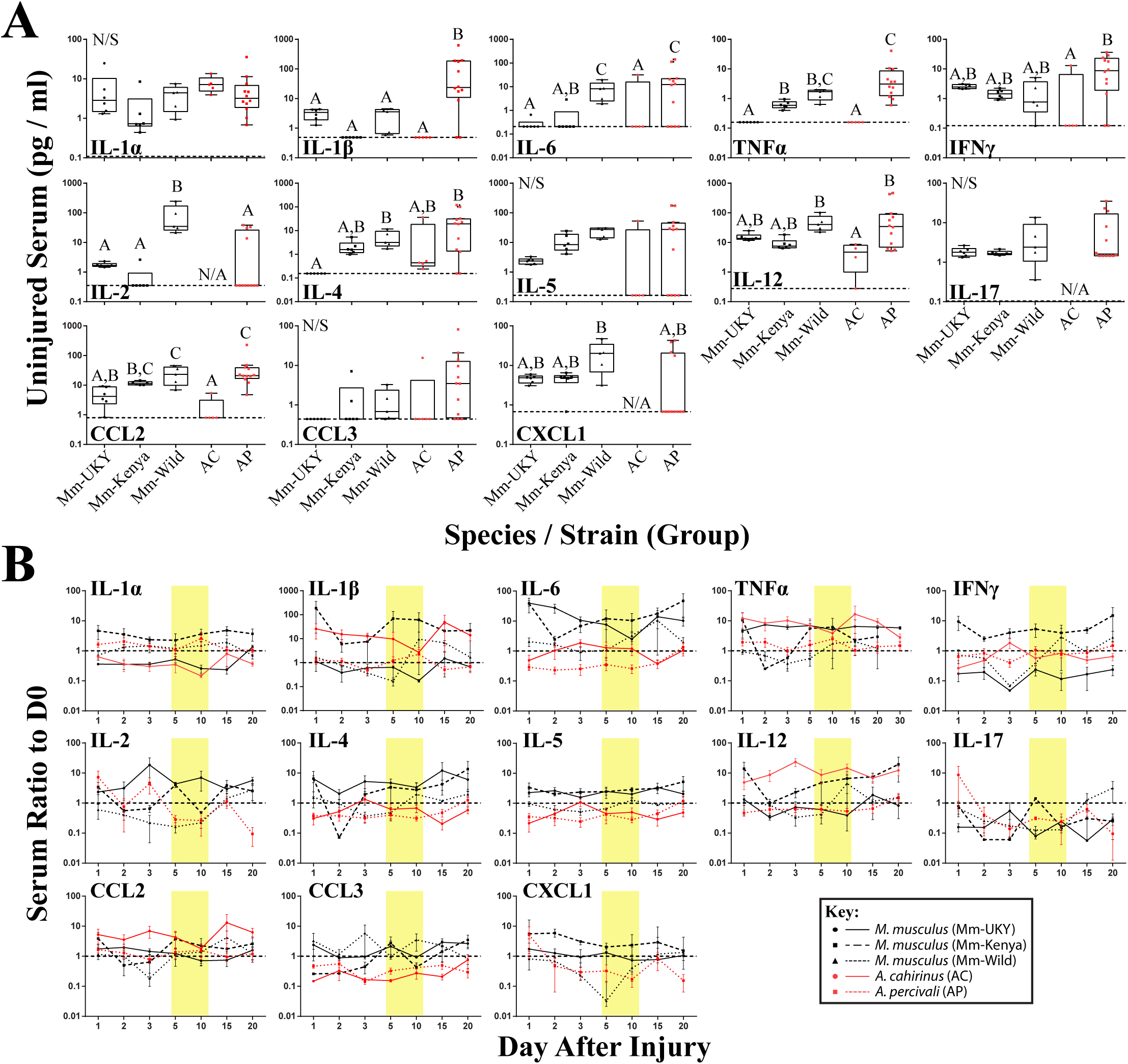
Ability to regenerate and immunity-status are associated with specific systemic immune profiles and immune responses to injury. **(A)** Comparison of cytokine concentrations in serum from uninjured animals showed higher concentrations of IL-4, IL-6, CCL2 and TNFα in wild-caught animals compared to laboratory-reared animals indicated that wild animals have a “primed” immune system. No difference was found between non-regenerators (Mm-UKY, Mm-Kenya and Mm-Wild) and regenerators (Ac and Ap). Data represent box and whiskers with median, interquartile range and individual data points. N/A denotes concentrations could not be quantified in any animal of the group. The dashed line in each graph represents the lower limit of detection for the specific cytokine. N/S denotes *P*>0.05 for One-way ANOVA (See Supplemental Table 1) and different letters above the data denotes *P*≤0.05 for Tukey-Kramer pairwise comparisons (See Supplemental File). **(B)** The change in systemic cytokines compared to D0 over 20 days after ear hole punch. A naïve immune system response (solid lines) was consistent with increased IL-2 and TNFα, and decreased IL-1α compared to a primed response (intermittent lines) (See Supplemental Table 2). Fibrotic repair (black) was associated with increases in IL-5, IL-6, IL-17, CCL3 and CXCL1 compared to regeneration (red). Data represent mean and S.E.M. for at least n=5 animals per species per timepoint. The dashed line at Y=1 represents no change compared to D0 and the yellow boxes represent the inflammation resolution window.

**Figure 2.**
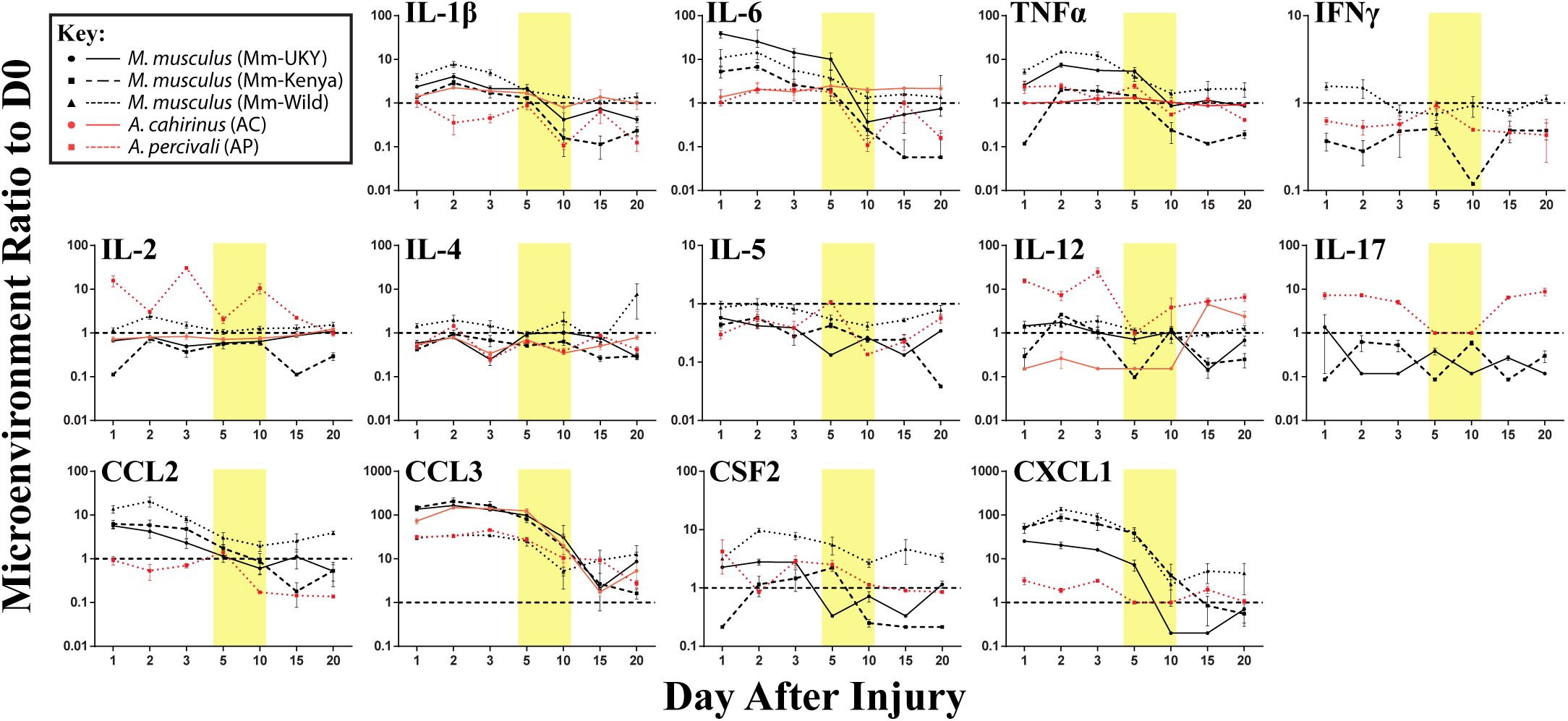
The tissue microenvironment is dynamic, and the ability to regenerate is associated with a specific immune response to injury. The comparison of the change in cytokines from tissue lysate compared to D0 over 20 days after ear hole punch. All cytokines varied over time and between groups (See Supplemental Table 3). There was a stronger increase in CCL3 a muted increase for IL-12 and CXCL1 in immune-naïve (solid lines) compared to immune-primed animals (intermittent lines). Non-regenerators (black) had stronger increases for IL-6, CCL2 and CXCL1 compared to regenerators (red). Additionally, IL-17 decreased in non-regenerators and increased in regenerators. Data represent mean and S.E.M. for at least n=5 animals per species per timepoint. The dashed line at Y=1 represents no change compared to D0 and the yellow boxes represent the inflammation resolution window.

Next, we quantified the systemic injury response for each cytokine compared to its baseline, beginning 24 hr (D1) after injury and over the next twenty days (Figure 1B). In most cases (except IL-2, IL-6, IL-17 and CXCL1), there was no effect of day (Supplemental Table 2), indicating that the immediate systemic response persisted for 20 days. Animals with a more naïve immune system showed increased IL-2 and TNFα, and decreased IL-1α compared to a relatively activated immune system (Figure 1B: solid lines compared to intermittent lines and Supplemental Data File). Regeneration showed decreased IL-5, IL-6, IL-17, CCL3 and CXCL1 compared to fibrotic repair (Figure 1B: red lines compared to black lines and Supplemental Data File). This latter result supported that animals healing by fibrotic repair and regeneration could be separated by their systemic response to injury.

### A regenerative microenvironment is marked by induction of T-cell-associated cytokines and a dampened pro-inflammatory cytokine response

Resident cells and infiltrating immune cells secrete cytokines that likely polarize the injury microenvironment to support regeneration or fibrotic repair [12, 13, 15, 16, 34, 46, 47]. Thus, to quantify local cytokine concentrations we assayed tissue lysate collected throughout the healing response. IL-1α could not be compared because baseline concentrations were above the upper limit of quantification in more than 80% of samples, indicating that IL-1α in the ear pinna was at least two orders of magnitude greater than the other cytokines measured. Although ear pinna tissue is structurally similar across species [40], local cytokines in *Acomys* were consistently detected at lower concentrations compared to *M. musculus*. Moreover, because the concentration of cytokine may not be as important as the dynamics of the cytokine, we compared the change in cytokine compared to baseline over 20 days (Figure 2). Injury elicited significant changes over time compared to baseline for all cytokines measured in tissue lysate supporting a dynamic response (Supplemental Table 3). Furthermore, while most cytokines shared similar trajectories over time, there was an effect of Group and the Group*Day interaction for all cytokines supporting significant differences in the magnitude of change among the groups (Figure 2 and Supplemental Table 3). Supporting an inflammatory response in all groups, several pro-inflammatory cytokines (IL-6, TNFα) and myeloid chemotactic factors (CCL3, CSF2 and CXCL1) showed an increase compared to baseline between D1 and D3 that then decreased to baseline or below between D5 and D20 (Figure 2). There was also an overall decrease compared to baseline for IL-5 and a small but significant change from baseline for IL-2 and IL-4 across all groups (Figure 2). Animals with naïve immune responses had a stronger increase in CCL3 and a smaller increase for CCL2 and CXCL1 compared to activated immune responses (Figure 2: solid compared to intermittent lines). We also identified several cytokines that showed differential changes between regeneration and fibrotic repair that we describe below (Figure 2: red compared to black lines).

During the acute inflammatory phase (D1 and D2), CCL2 and CXCL1 were increased 9 and 12-fold during fibrotic repair compared to regeneration, respectively (Figure 3). IL-6 showed a similar result at D2 where Mm-UKY, Mm-Kenya and Mm-Wild were increased 10-fold compared to Ac and Ap (Figure 3). Additionally, IL-17 was increased in Ap, but decreased in Mm-UKY and Mm-Kenya and IL-12 was increased in Ap compared to all *Mus* (Figure 3). Interestingly, the TNFα response—a biomarker of inflammation—could not reliably separate fibrotic repair and regeneration (Figure 3).

**Figure 3.**
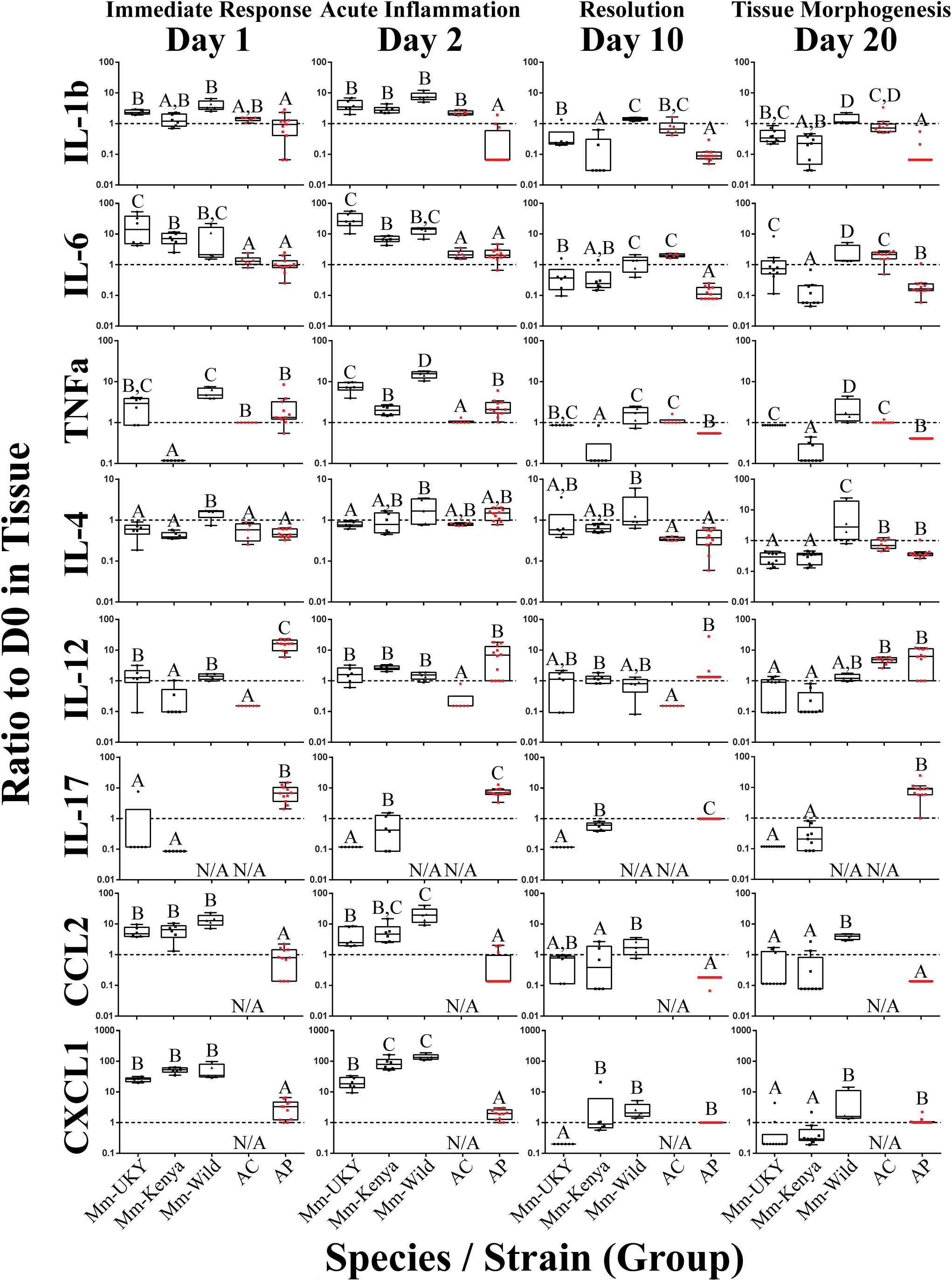
Regeneration is associated with specific temporal cytokine responses in the tissue microenvironment. The comparison of the change in cytokines from tissue lysate compared to baseline. Non-regenerators (Mm-UKY, Mm-Kenya, and Mm-Wild: Black) show a stronger response compared to regenerators (Ac and Ap: Red) for several inflammatory cytokines during the immediate response, acute inflammatory phase. These differences mostly resolved at day 10 and 20. In contrast, regenerators showed a stronger IL-12 and IL-17 response during tissue morphogenesis. Data represent box and whiskers with median, interquartile range and individual data points. N/A denotes concentrations could not be quantified. The dashed line at Y=1 represents no change compared to D0. Each graph showed P<0.05 for effect of group using a One-way ANOVA on log-transformed data (See Supplemental Table 4) and different letters above each group denotes *P*<0.05 for Tukey-Kramer pairwise comparisons (See Supplemental File).

Regardless of healing outcome, re-epithelialization occurs by D10 [40, 42] coincident with resolution of many pro-inflammatory cytokine responses (Figure 1B and 2: yellow bars). While there were some differences among groups for the timing of resolution, IL-1β, TNFα, and CCL2 were similar to or below baseline at D10 for each species (Figure 3). IL-6 also followed this pattern; however, there was a differential response where Ac remained elevated through D20 while all other species decreased below baseline (Figure 3).

At D20, during tissue morphogenesis, the only cytokines that showed a differential response were IL-12 and IL-17 that were increased during regeneration compared to fibrotic repair (Figure 3). The anti-inflammatory cytokine IL-4 did not differ over time with respect to regenerative ability, suggesting that the differences in pro-inflammatory cytokine release is likely not an IL-4 mediated response. Our results suggest that subtle differences in how cytokines are deployed in the injury microenvironment can distinguish regeneration or fibrotic repair. These data suggest that strong, acute increases in the pro-inflammatory cytokines IL-6, CCL2 and CXCL1 are associated with fibrosis, while the release of IL-12 and IL-17 during tissue morphogenesis is associated with regeneration.

### Regeneration is associated with an early burst of T cell influx to the injury site

The release of IL-12 and IL-17 into the regenerative microenvironment suggested enhanced T cell activation during regeneration [48, 49]. Therefore, we quantified T cell influx into uninjured and healing tissue from our laboratory populations of *Mus* (Mm-UKY) and *Acomys* (Ac) using flow cytometry with an antibody to the extracellular portion of the T cell marker CD3. We observed significant differences in CD3+ cells in injured tissue between species over time (two-way ANOVA, n=57; species: Df=1, F=49.49 *P*<0.001; day: Df=6, F=89.07, *P*<0.001; species*day: Df=6, F=21.49 *P* < 0.001) (Figure 4A). In uninjured tissue, Mm-UKY had 10-times more CD3+ cells compared to Ac (Tukey-Kramer HSD post-hoc test, Df=6, t=6.21, *P*<0.001) (Figure 4A). While the total number of CD3+ cells that infiltrated the wound was higher in Mm-UKY compared to Ac, there was a greater fold change relative to D0 for CD3+ cells during regeneration compared to fibrotic repair (Figure 4B). Ac exhibited a monophasic response to injury starting on D1 with a 78-fold influx of T cells that peaked on D3 and remained above baseline at D15. Mm-UKY showed a biphasic response with peak influx of 10-fold at D7 that returned to baseline at D15 (Figure 4B). Importantly, at D15, when IL-12 was increased (Figure 2), the influx of CD3+ cells remained high in Ac compared to Mm-UKY (Figure 4B).

**Figure 4.**
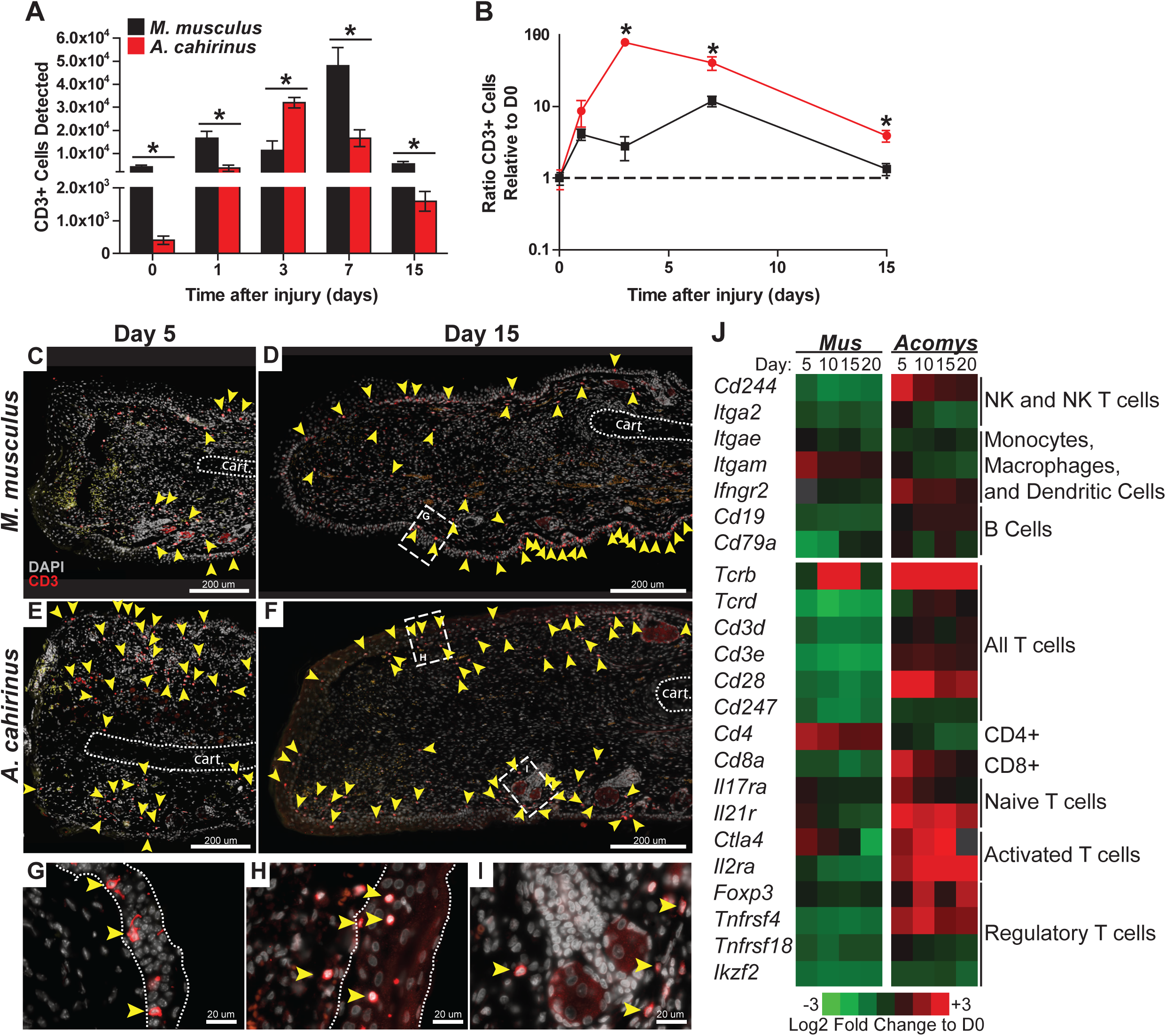
The regeneration microenvironment is primed by greater T cell influx and activated TREG signature. **(A)** Comparison of total CD3+ cells quantified by flow cytometry from disassociated ear pinna and **(B)** the ratio relative to uninjured tissue for *M. musculus* (black) and *A. cahirinus* (red). Data represent mean and S.E.M. and n=4 or 5. An * denotes *P*<0.05 for pairwise comparison within the day between species for Tukey-Kramer HSD posthoc test. **(C-I)** Representative immunohistochemistry for CD3 (red) counterstained with DAPI (gray) at the proximal wound margin (amputation plane can be determined from the end of the cartilage— indicated by the dotted line) from D5 and D15 after injury of *M. musculus* (C, D) and *A. cahirinus* (E, F). More T cells (yellow arrowhead) were present throughout the wound bed and were mainly found in the dermis of *A. cahirinus* compared to *M. musculus*. The T cells associated with epidermis (boundaries indicated by the dotted line) tended to be spindle-like in *M. musculus* (G), while rounded in *A. cahirinus* (H). The dermal T cells in *A. cahirinus* also tended to be in close proximity to regenerating epidermal appendages (I). N=4 and bar equals 200 µm (C-F) or 20 µm (G-I). **(J)** Heatmap of differential gene expression compared to uninjured tissue suggests that the regeneration microenvironment contains a substantial NK, CD8+ and T_REG_ cell response while fibrotic repair has a CD4+ cell response. Data comes from a previously published analysis [40].

We next used immunohistochemistry with an antibody specific to the intracellular portion of the CD3 receptor to assess the spatial distribution of T cells during acute inflammation and morphogenesis (Figure 4C-E). First, these data confirm the greater influx of CD3+ cells at D5 and D15 in Ac compared to Mm-UKY (Figure 4C-E). Second, in Mm-UKY most CD3+ cells were associated with the epidermis and were rarely observed distal to the amputation plane (Figure 4C). On the other hand, CD3+ cells in Ac were present in the epidermis and dermis, and regularly observed in healing tissue distal to the amputation plane (Figure 4E). At D15, CD3+ cells were found in the epidermis and dermis of both species (Figure 4D, F). Interestingly, CD3+ cells associated with the epidermis in Mm-UKY (Figure 4G) exhibited a spindle-shape morphology compared to a rounded shape in Ac (Figure 4H). There also appeared to be more CD3+ cells in the dermis of Ac compared to Mm-UKY (Figure 4D, F), and the CD3+ cells tended to localize near regenerating hair follicles in Ac (Figure 4I). Attempts to characterize individual T cell phenotypes during regeneration using flow cytometry and IHC using 19 commercially available antibodies, supported significant differences in antibody-epitope binding between species that prevented further T cell phenotyping by receptor subtype in *Acomys* (Supplemental Table 5). Therefore, we interrogated a comparative injury RNAseq dataset for differential expression of T cell associated transcripts between *Mus* and *Acomys* [40]. While expression for genes associated with non-lymphocyte immune cell populations were generally similar between species, several transcripts associated with T cells and natural killer cells were increased in *Acomys* and decreased in *Mus* in response to injury (Figure 4J). Increased expression of *Cd8*, *Ctla4*, *Il2ra*, *Foxp3*, and *Tnfrsf4* specifically suggested an activated cytotoxic and regulatory T cell response during regeneration but not fibrotic repair (Figure 4J). During fibrotic repair, *Cd4* was differentially increased at D5 and D10 suggesting the presence of CD4 helper T cells not present during regeneration (Figure 4J). Together, these data demonstrate that regeneration was associated with a proportionally greater influx of CD3+ cells that accumulate quickly at the injury site and that specific subtypes of activated T cells were preferentially associated with regeneration.

### STAT3 is activated independently from IL-6 during blastema formation

We also sought to test our observation that strong induction of the pro-inflammatory cytokine IL-6 was associated with the acute inflammatory phase of fibrotic repair. To do this, we assayed for IL-6 signaling using STAT3 phosphorylation (Figure 5A-F). STAT3 is phosphorylated in response to the ligand IL-6 binding its membrane receptor, which activates signal transduction in target cells [50]. Corroborating our ELISA quantification for IL-6 in the tissue microenvironment, we found that pSTAT3 increased 8-fold in response to injury in Mm-UKY during the acute inflammatory phase (Figure 5A, B). Similarly, during fibrosis when IL-6 concentrations resolved in Mm-UKY, pSTAT3 began to decline toward baseline (Figure 5A, B). In *Acomys*, pSTAT3 was significantly elevated at D1, although to a lesser extent than compared to Mm-UKY (Figure 5A, B). Moreover, during blastema formation (D10-15) when our ELISA data showed increased IL-6 compared to baseline in Ac (Figure 3), analysis of pSTAT3 showed further induction of pSTAT3 in Ac (Figure 5A, B).

**Figure 5.**
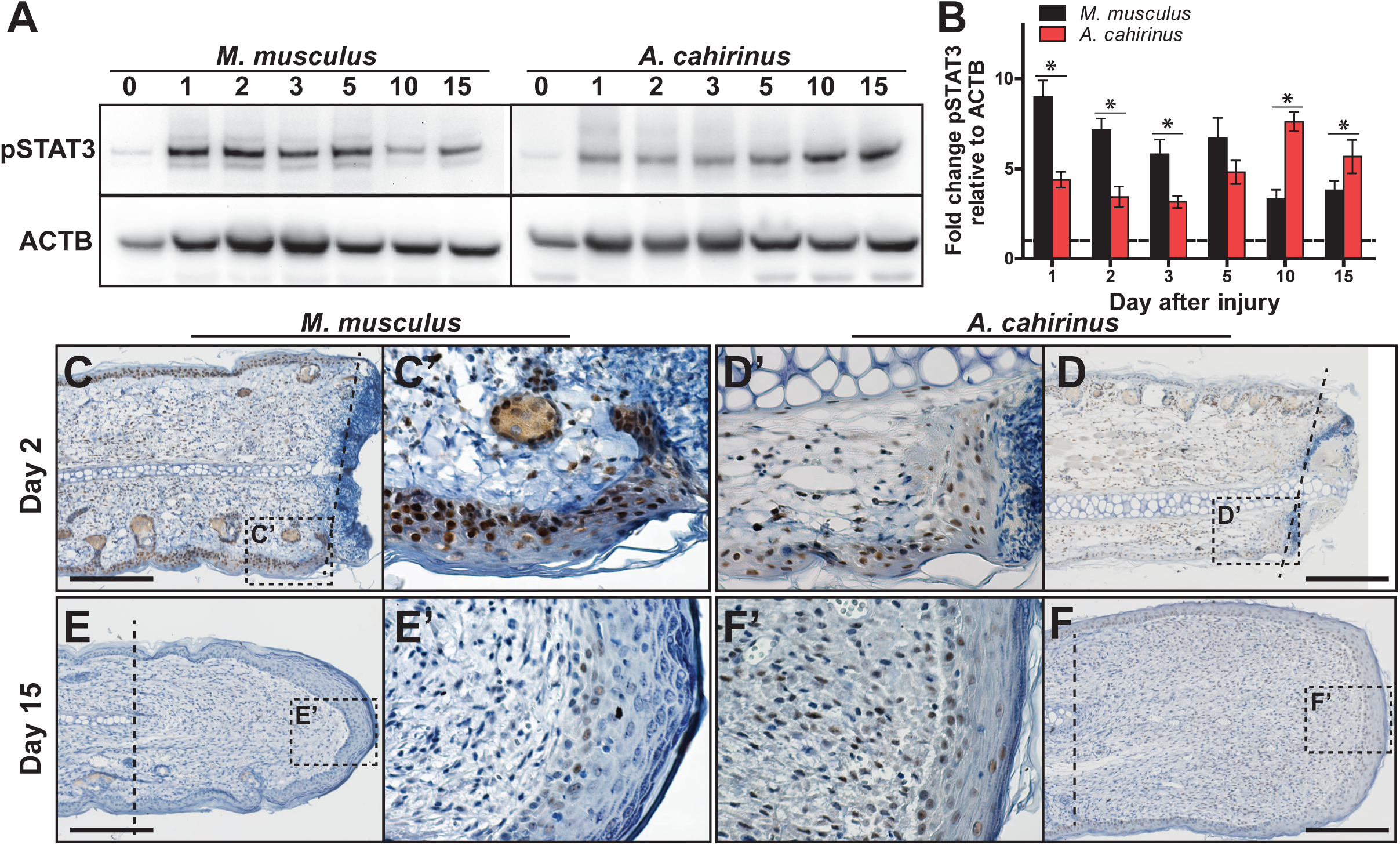
Time-dependent STAT3 activation in the blastema is associated with regeneration. **(A-B)** Comparison of representative immunoblots for pSTAT3 and ACTB for indicated time and species from injured tissue homogenate from ear-hole punch assay (A). Fibrotic repair is associated with strong early STAT3 activation while regeneration is associated with a weak early and strong postponed activation (B). Data represent mean fold change of pSTAT3 intensity normalized to ACTB intensity and relative to D0 and S.E.M. n=4 individuals per time point and species (B). (**C-F**) Representative images for immunohistochemistry for pSTAT3 (Brown) counterstained with hematoxylin (Blue) for *M. musculus* (C, C’, E and E’) and *A. cahirinus* (D, D’, F and F’) at the indicated time points. On D2 after injury, nearly every epidermal cell in *M. musculus* was positive compared to about half in *A. cahirinus* (C, D). On D15 after injury, only a small population of epidermal cells in *M. musculus* were positive compared to about half in *A. cahirinus* (E, F). Additionally, about half of the cells in the forming blastema remain positive at D15. Data represent n=3, and bar equals 200 µm.

To determine the cellular localization of STAT3 phosphorylation, we assayed for pSTAT3 using immunohistochemistry during the acute inflammatory phase (D2) and new tissue formation (D15) (Figure 5C-F). Supporting the immunoblot data, both species showed extensive nuclear staining for pSTAT3 at D2 in the epidermis and mesenchymal compartments (Figure 5C’, D’). Positive staining in the epidermis > 200 µM proximal to the amputation plane suggested STAT3 activation was a pervasive response to injury within the ear pinna in both species (Figure 5C, D). Supporting the 2-fold difference in pSTAT3 we observed between Mm-UKY and Ac (Figure 5A, B), we found that nearly every epidermal cell in Mm-UKY appeared positive for pSTAT3 whereas less than half of the epidermal cells were positive in Ac (Figure 5C’, D’). The internal tissue compartments (e.g., dermis, cartilage, muscle and adipose) at D2 were similar between species with approximately half of the total cells positive for pSTAT3. At D15, only a few pSTAT3 positive cells were present in Mm-UKY and they were isolated to the epidermis distal to the amputation plane (Figure 5E, E’). In contrast, pSTAT3 positive cells were widespread throughout the blastema in Ac (Figure 5F, F’). Together, these data support stronger IL-6 mediated STAT3 activation in Mm-UKY compared to Ac during the acute inflammatory phase and increased STAT3 activation during blastema formation.

Greater increases in IL-6 and CXCL1 during the acute inflammatory phase of fibrotic repair in *M. musculus* suggested that these molecules might antagonize a potential regenerative response. Previous studies have shown that a balance in these molecules regulate wound healing as IL-6 and CXCL1 are potent pro-inflammatory molecules and hyper-elevated concentrations after injury are attributed to aberrant healing and chronic inflammation [51–53]. Additionally, genetic ablation of *IL-6*, the *IL-6 receptor*, or the *CXCL1 receptor* (*CXCR2*), causes severely delayed re-epithelialization, scab formation and abhorrent wound healing in cutaneous and incisional wounds [54–57]. IL-6 signaling activates several downstream mediators of inflammation including cyclooxegenase-2 (COX-2), and its enzymatic products can amplify the inflammatory response [58]. To test if COX-2 activity promotes fibrosis in the ear pinna, we used our ear punch assay in Mm-UKY treated healing tissue with Celecoxib, a specific and potent COX-2 inhibitor [59] (Figure 6A). Comparing ear-hole closure between celecoxib- and vehicle-treated animals there was no support for a difference in the rate of closure (D5 through D30) (repeated measures two-way ANOVA, n=31; treatment: Df=1, F-2.42, *P*=0.137; day: Df=5, F= 179.62, *P*<0.001; treatment*day: Df=5, F=0.36, *P*=0.875) (Figure 6B). Similarly, comparison of ear-hole area at D64 showed no support for a difference between treatment and control ears (unpaired t-test; t=0.671, *P*=0.512) (Figure 6C). One out of five celecoxib-treated animals had not completed re-epithelialization by D10 (Figure 6D-F). Lastly, while the intensity of Picrosirius stain appeared to be lower in celecoxib treated animals compared to controls, there was no difference in the area of collagen deposition at D64 (unpaired t-test, t=0.104, *P*=0.918) (Figure 6G-I). Thus, these data support that systemic inhibition of COX-2 activity is not sufficient to reduce fibrosis or induce a regenerative response after ear pinna injury in *M. musculus*.

**Figure 6.**
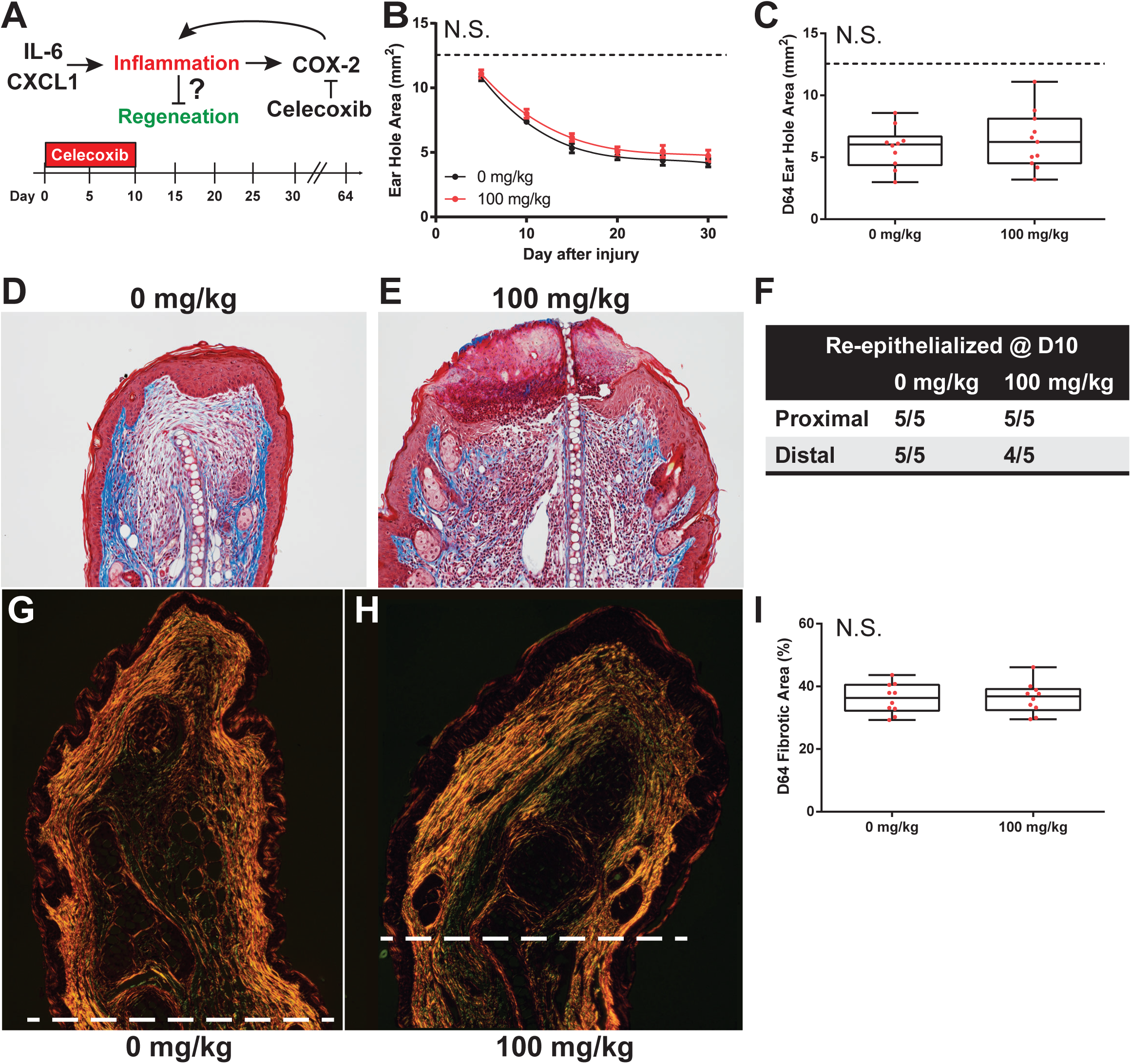
COX-2 inhibition does not liberate regeneration in *M. musculus*. **(A)** Celecoxib treatment during the first 10 days of injury reduces secondary inflammation caused by IL-6 and CXCL1. **(B-C)** Celecoxib treatment did not affect the rate of ear-hole closure from D5 to D30 (B), or ear-hole area at D64 (C). **(D-F)** Mason’s trichrome stain on D10 ear tissue from control (D) and treatment (E) showed 20% of the distal injury plane from treated ears were not re-epithelialized, while 100% of control treated ears and the proximal injury plane were re-epithelialized (F). **(G-I)** Picrosirius stain on D64 ear tissue from control (G) and treatment (H) showed no difference in the total collagen fibrotic area distal to the amputation plane (I). Data represent mean +-S.E.M. and the lines represent cubic regression for n=10 per treatment (B), OR indivdual data (red dots) and median and interquartile range for n=10 per treatment (C, I).

## DISCUSSION

In this study, we performed a comprehensive cytokine characterization of the immune response to injury where identical injuries in closely-related species undergo two different healing responses: regeneration or fibrotic repair. Importantly, our experimental design leveraged a comparison of animals with an activated and naïve immune system in order to identify species-specific cytokine changes that were associated with regeneration and not due to an environment-immunity interaction. Our analyses showed that regardless of healing outcome, injury induced a common set of pro-inflammatory factors (IL-6, and TNFα) and chemokines (CCL3, CSF2 and CXCL1) during the acute inflammatory phase of fibrosis and regeneration. Additionally, we observed similar responses between healing outcomes for IL-2, IL-4 and IL-5 in the tissue microenvironment. While this supports that fibrotic repair and regeneration share inflammation during the healing process, we also found significantly greater responses for IL-6, CCL2 and CXCL1 during fibrotic repair compared to regeneration. In contrast, regeneration was uniquely associated with local increases in IL-12 and IL-17. Supporting our cytokine analysis, we found that regeneration was associated with a strong influx of T cells during acute inflammation compared to fibrotic repair and that regeneration-competent T cells were closely associated with the dermis during blastema formation. This latter point suggests that T cells may influence local inflammation and contribute to the local regulation of tissue morphogenesis in spiny mice.

Recent studies comparing immune profiles between laboratory-reared and pet-store or wild-caught *M. musculus* demonstrate that non-laboratory strains have more CD44^+^ effector T cells, memory T cells and circulating neutrophils [60, 61]. Although neither group directly measured serum cytokines from the different populations, the elevated baseline concentrations of IL-4, IL-6, CCL2 and TNFα that we measured in circulation from immune-challenged animals support larger active populations of effector and memory T cells. These data support that our immune-challenged group have been exposed to more pathogens than the laboratory-reared mice which is undoubtedly the case. In addition to the increased baseline concentrations of these cytokines, we also found significant differences in the response to injury for IL-1α, IL-2 and TNFα between animals with an activated or naïve immune system. Studying wild-caught populations enabled us to identify responses that accurately reflected phenotypic differences between species, rather than differences that could be explained by immune status. Of particular importance was our inclusion of wild-caught *A. percivali* that indicate increases in TNFα, CCL2 and CXCL1 are not inhibitory to regeneration. Additionally, we observed high variation in cytokine concentrations across our dataset, indicating that the immune response to injury could be confounded by individual variation. In other words, researchers should not expect to identify clear transition phases based on time after injury with small sample sizes. Ultimately these data support that injury elicits a local cytokine response that is dependent of baseline immune status with respect to release of cytokines that in turn effects the timing of events but does not change healing outcome.

Acute inflammation is a necessary component of the innate immune response designed to fight invading microbes by recruiting leukocytes from circulation and activating local myeloid and lymphoid cells. Our analyses demonstrate that injury induces an acute inflammatory response regardless of healing outcome that resolves within ∼10 days; a timeframe in line with human and rodent wound healing studies [7, 62]. In particular, we found the local release of CCL3, CSF2 and CXCL1 in all groups, which are known to be potent chemokines for monocytes, macrophages and neutrophils. Moreover, the local release of IL-6 and TNFα supports the presence of activated macrophages and neutrophils as a common injury response. Supporting our previous work that neutrophils infiltrate injured spiny mouse tissue slower compared to laboratory mice [15], regeneration was associated with delayed release and a reduced maximal fold change in IL-6 and CXCL1 compared to fibrotic repair. Additionally, we did not detect a CCL2 response during regeneration, and IL-6, CXCL1 and CCL2 are known to positively regulate the speed of re-epithelialization [54–56], which we find to be delayed at most five days in *A. cahirinus* compared to *M. musculus* [40]. Thus, while acute inflammation is a component of regeneration and fibrotic repair, the cellular differences are likely attributed to reduced pro-inflammatory cytokines released into the microenvironment.

Corroborating our observation that the IL-6 response was weaker during regeneration compared to fibrotic repair, we found diminished activation of STAT3 during acute inflammation (D1-10) in spiny mouse epidermis compared to mouse. Interestingly, we observed an increase in pSTAT3 during blastema formation, whereas pSTAT3 levels declined during fibrotic repair. Furthermore, during tissue morphogenesis at D15 many blastemal cells were STAT3 positive. Given that IL-6 concentrations did not appreciably increase during blastema formation or tissue morphogenesis the increase in STAT3 activity is likely independent of IL-6. STAT3 is activated through multiple pathways (e.g. leukemia inhibitory factor, epidermal growth factor, palette derived growth factor, IL-10, IL-17, etc.). Although IL-17 increased in *A. percivali* after D10, it did not increase in *A. cahirinus* suggesting it is not responsible for the late phase of STAT3 phosphorylation. Given that STAT3 signaling is multifaceted, one potential biological link is that STAT3 activity is necessary for satellite-cell activation and axon regeneration in mammals [63–65]. Interestingly, the expression of *Sal4—*a factor necessary for blastema maintenance in *Xenopus* and *Ambystoma—*is regulated by pSTAT3 [66–68]. While *Sal4* does not have a mammalian homolog, this data supports that activation of STAT3 in regenerating tissue is an evolutionary conserved mechanism and interrogating unique STAT3 targets in spiny mice may uncover mechanisms that regulate blastema formation in mammals.

Inhibition of downstream signaling induced by IL-6 / CXCL1, such as arachidonic acid metabolism by COX-2, has been shown to reduce fibrosis post epidermal injury (e.g. incisional, cutaneous and chronic pressure wounds) [69–71]. Celecoxib treatment to inhibit COX-2 in the present study may have slowed re-epithelialization. Additionally, while the total area of fibrosis was not different between celecoxib- and vehicle-treated animals there appeared to be a small reduction in the total amount of collagen produced in celecoxib-treated animals from reduced intensity of picrosirius staining. However, similar to previous reports, reduction in COX-2 activity did not induce regeneration, supporting that inflammation is not the ultimate inhibitory barrier.

In addition to the magnitude increase in IL-6 and CXCL1, our analyses found that increased local CCL2 was specific to fibrotic repair. CCL2 was first identified as a monocyte-specific chemoattractant to sites of injury and infection, and activates macrophages [72, 73]. CCL2 also attracts neutrophils and supports neutrophil-dependent tissue damage [72]. As such, the amount of CCL2 that is released into an injury microenvironment regulates the healing response and studies support there is a positive relationship between CCL2 and the amount of fibrosis during fibrotic repair [74–76]. However, a careful balance must be maintained as CCL2 knockout mice do not heal wounds [77]. Thus, it is possible that the reduced IL-6, CCL2 and CXCL1 responses are responsible for reduced fibrosis in spiny mice. Although these key factors appear to interact in the hierarchy of the progression of fibrotic repair, the paracrine mechanism of how they would activate dermal fibroblasts remains unknown. It is likely another cell-type, such as a macrophage or T cell, is mediating the signal.

In addition to this study, two studies have quantified cytokines during regeneration—one in axolotl limbs [12] and the other in spiny mouse dorsal skin wounds [34]. Godwin et al. (2013) used a mouse cytokine array to analyze regenerating salamander limbs and found that all but two cytokines detected reached peak amounts within 48hrs of injury and that every cytokine returned to baseline by D15 after a blastema had formed. Brant et al. (2016) used the same cytokine array to assess cytokines during the first 14 days of spiny mouse dorsal skin regeneration and observed a similar phenomenon and all detected cytokines resolve to baseline by D14. Despite study-specific differences in the ability to detect antigens and a lack of parallelism validation, these studies support that release of CCL3 and TNFα in tandem with a differential inflammatory response occurs prior to tissue regeneration. However, our comparative analyses also suggested that the magnitude of the increase in IL-6 and CXCL1 might serve as early indicators of a fibrotic repair trajectory. For example, the IL-6 response to injury, although present, was small and CXCL1 did not respond during both axolotl limb and spiny mouse skin regeneration.

Finally, our cellular analysis uncovered a surprisingly rapid adaptive immune response measured as an early influx of T cells in regenerating compared to non-regenerating species. Importantly, our findings support that the arrival of T cells in spiny mice is concurrent with the arrival and proliferation of monocytes [15], which suggests there is a regenerative-competent T cell response that is different from a fibrotic T cell response. Contrary to hypotheses suggesting that a strong adaptive immune response reduces regenerative ability [78, 79], our findings suggest T cells may positively regulate regeneration in spiny mice. Similar to macrophages, T cells can differentiate into a number of functional subpopulations which differentially affect cells in the inflammatory microenvironment [80]. Our analysis of the transcriptional response to injury between *M. musculus* and *A. cahirinus* suggests that fibrotic repair is associated with an accumulation of inactivated T_H_ cells, while during regeneration there is an accumulation of activated cytotoxic T and regulatory T (T_REG_) cells. Studies have shown that loss of cytotoxic CD8^+^ T cells inhibits skeletal muscle regeneration, accelerates bone fracture healing, and increases fibrosis in incisional wounds [81–87]. Additionally, recent work showed that T_REG_ populations infiltrate injured muscle quickly after injury and are necessary to regulate the ratio of MHC-class II positive and negative macrophages present in the injured tissue. When the T_REG_ population is ablated subsequent regeneration is impaired [17]. Moreover, spiny mice have a greater NADPH oxidase induced ROS response [15], which can be partially controlled by T_REG_ cells [88–90]. Thus, these studies support an anti-fibrotic role for T cells and suggest the action of cytotoxic T and T_REG_ cells may have a positive role during spiny mouse epimorphic regeneration.

Together, the data presented here support that tissue regeneration in *Acomys* occurs in cooperation with an adaptive immune response and that lymphocyte phenotype might play a key role in facilitating a regenerative or fibrotic response. Ongoing studies in our laboratory are aimed at characterizing the macrophage and T cell populations that are associated with the injured tissue during regeneration and fibrotic repair. These future datasets (and the present one) will create a framework to begin testing how the immune response functions during complex tissue regeneration in a mammalian model. We believe that modulating the immune response at the injury microenvironment will be an essential piece to inducing epimorphic regeneration in tissues that naturally heal by fibrotic repair.

## AUTHOR CONTRIBUTIONS

Conceptualization, T.R.G., V.O.E. S.G.K. and A.W.S.; Methodology, T.R.G., J.S., and A.W.S; Validation, T.R.G. and J.S.; Investigation, T.R.G, J.S., J.M.K., C.K.H., V.O.E. and A.W.S.; Resources, S.G.K., V.O.E. and A.W.S.; Writing – Original Draft, T.R.G.; Writing – Review & Editing, T.R.G., J.S., V.O.E. and A.W.S.; Visualization, T.R.G. and A.W.S.; Supervision, T.R.G., S.G.K., V.O.E. and A.W.S., Project Administration, T.R.G., V.O.E. and A.W.S.; Funding Acquisition, S.G.K., V.O.E. and A.W.S.

## ACKNOWLEDGEMENTS

We thank Adam Cook, Chanung Wang, Bailey Gensheimer, Malik Guidry, John Ewoi and Stanley Marete for help with live-trapping, animal care and data collection. We also thank Peter Jessel and the Tomlinson family for kindly allowing us continued access to their property in Kenya. We acknowledge David Higginbotham, Jim Monegue and rest of the swine unit for their help and enthusiasm with trapping the wild *Mus musculus*. Funding for this work was provided by the National Science Foundation (NSF) and the Office for International Science and Engineering (OISE) to A.W.S. (IOS-1353713) and V.O.E (IOS-1353857). The University of Kentucky provided additional funding to A.W.S.

## MATERIALS AND METHODS

### Animals

*Mus musculus* (Mm-UKY) and *Acomys cahirinus* (Ac) were maintained at our animal facility at the University of Kentucky, Lexington, KY, USA. Mm-UKY were sexually mature (10- to 12-week old), female, outbred Swiss Webster (ND4, Envigo, Indianapolis, IN). They were housed at a density of 2-4 individuals in static IVC cages with pine shavings and given autoclaved water and 18% protein mouse chow (Tekland Global 2918, Envigo). Ac were sexually mature (12- to 28-weeks old), males and females and were housed at a density of 10-15 individuals in large metal wire cages (24 inch x 12 inch x 14 inch, height x width x depth, Quality Cage Company, Portland, OR) with pelleted pine bedding (Southern States Cooperative, Inc., Richmond, VA) and given autoclaved water and a 3:1 mixture by volume of 14% protein mouse chow (Teklad Global 2014, Envigo) and black-oil sunflower seeds (Pennington Seed Inc., Madison, GA) [91]. Additionally, the air within facility was filtered, and the animals were exposed to natural light through large windows (approximately 12h:12h L:D light cycle during the experiment). All Mm-UKY and Ac samples were collected between 9/20/2015 and 10/28/2015.

Wild *M. musculus* (Mm-Wild) were live trapped at the C. Oran Little Research Center in Versailles, KY (38°4’N, 84°44’W) and maintained in an alternate animal facility at the University of Kentucky. Mm-Wild were housed at a density of 10-12 individuals in large metal wire cages with pelleted pine bedding and given autoclaved water and 18% protein mouse chow. The animals acclimated to captivity for at least twenty-one days before any experiments were started. The air within the facility was filtered and the animals were exposed to a 12:12h L:D cycle by fluorescent lights. All Mm-Wild samples were collected between 4/1/2017 and 6/21/2017 and 12/12/2017 and 3/6/2018.

The Kenyan *M. musculus* (Mm-Kenya) were sexually mature (10- to 12-week old), female, outbred Swiss Webster mice obtained from a local breeder in Nairobi, Kenya and maintained in an animal facility at the University of Nairobi, Kenya. Sexually mature *Acomys percivali* (Ap) were live-captured at Mpala Research Centre in Laikipia, Kenya (0°17’N, 37°52’E), and transported to the University of Nairobi for study. Each species was separated by sex and housed at a density of 10-15 animals in large metal wire cages, given tap water, fed mouse pencils (Argrocide Inc., Nairobi, Kenya) 1x per day and exposed to natural light through windows (equivalent 12h:12h L:D cycle). The animals acclimated to captivity for at least twenty-one days before any experiments were started. Additionally, the facility was open to the natural environment (i.e., the mice were exposed to Nairobi air) and cool nighttime temperatures were supplemented with ceramic heaters. Mm-Kenya samples were collected between 6/04/2015 and 7/04/2015 and Ap samples were collected between 5/04/2015 and 7/04/2015, and between 5/02/2016 and 6/04/2016.

All animal trapping and procedures were approved by the University of Kentucky Institutional Animal Care and Use Committee (IACUC) under protocol 2013-1119, Kenyan Wildlife Service (KWS), and the University of Nairobi Faculty of Veterinary Medicine Animal Care and Use Committee (FVM ACUC). Research in Kenya was approved by the Kenyan National Council for Science and Technology (NACOSTI). All wild species trapped were species of least concern.

### Sample collection and preparation

We used a 4 mm biopsy punch to create a hole through the ear pinna, as previously described [40]. Healing ear tissue was collected on D0, 1, 2, 3, 5, 10, 15 and 20. To minimize circadian effects, animals were injured between 10:00 and 12:00, and samples were collected between 11:00 and 15:00. Animals were deeply anesthetized with 5% (v/v) isoflurane and a maximal amount of blood was collected by cardiac puncture using a 25-guage needle. An 8 mm biopsy punch was used to harvest healing ear tissue.

To isolate serum, blood was collected into a serum separator tube (#454243, Greiner bio-one, Kremsmünster, Austria) and allowed to clot for at least 45 minutes, followed by centrifuging at 3,000 x *g* for 10 minutes. Serum was aliquoted and stored at −80°C or on dry ice until analysis. The tissue was used for two downstream assays, histology and cytokine quantification. For histology, one of the 8 mm biopsies was placed into 10% (v/v) neutral buffered formalin (American MasterTech, McKinney, TX) overnight, dehydrated, embedded in paraffin and cut to 5 µM thickness on a rotary microtome. For cytokine quantification, a ring of tissue closest to the injury approximately 1 mm wide was snap frozen in liquid nitrogen or a slurry of dry ice and ethanol, and then stored at −80°C or on dry ice. Next, the tissue was homogenized in RIPA buffer supplemented with protease and phosphatase inhibitors (#24948, Santa Cruz Biotechnology, Inc. Dallas, TX; #78427, Thermo Scientific) using ceramic beads (Matrix D, MP Biomedicals, LLC, Solon, OH) and a bead mill for 5 minutes (Next Advance, Inc., Troy, NY), centrifuged at 10,000 x *g* for 15 minutes to pellet insoluble protein, and the soluble protein was separated into a new tube. The total protein was quantified by bicinchoninic acid assay (#23225, Thermo Scientific) with a standard curve created from the same stock of bovine serum albumin, and then the protein lysate was stored at −80°C or on dry ice until analysis.

### Cytokine assay

To assess the immune response to injury in multiple species, we evaluated methods that: 1) used minimal sample, 2) measured local (tissue lysate) and systemic (serum) samples, 3) measured several cytokines at once, 4) differentiated the magnitude and type of immune response during an ear punch assay, and 5) exhibited cross-reactivity among the study species. We used a custom-designed, multiplexed, sandwich ELISA array (Quansys Biosciences, Logan UT). This platform meets the above requirements and the experiments can be performed in multiple locations (i.e., Kentucky and Kenya) because the imager and reagents can be easily transported. Importantly, the imager does not require specialized calibration after being moved, and the reagents do not need remain frozen. The custom assay was designed to measure 16 antigens including IL-1α, IL-1β, IL-2, IL-4, IL-5, IL-6, IL-10, IL-12p70, IL-17, CCL2, CCL3, CCL5, CSF2, IFNγ, TNFα and CXCL1.

Initial testing identified that antigens from serum could be quantified by diluting serum in the supplied mouse specific diluent 1:1 and from tissue lysate using 5, 40, and 80 µg total protein in RIPA buffer for *M. musculus*, *A. cahirinus*, and *A. percivali,* respectively. The samples were run in duplicate using a protocol modified from the manufacturer’s instructions, as follows: All serum samples were diluted 1:1 (serum : diluent) and all tissue samples were diluted 1:1 (RIPA + lysate : mouse sample diluent) to a volume of 50 µL per well. The diluted samples were then loaded onto a new assay plate with an appropriate standard curve (1:3 to 1:59049) and four blanks. Samples were incubated at 4°C for 8 hours on a plate shaker set to 500 rpm to capture antigen in each well. After washing the plate 4 times with wash buffer, the primary antibody cocktail was loaded and the plate was incubated at 4°C for 8 hours on a plate shaker set to 500 rpm to allow binding of the biotinylated detection antibodies to the captured antigens. After washing the plate 4 times, streptavidin-HRP conjugated secondary antibody was loaded and the plate was incubated at room temperature for 30 minutes on a plate shaker set to 500 rpm. The plate was washed 8 times, chemiluminescent reagent was added, and the plate was immediately imaged with a chemiluminescent plate imager set to the manufacturer recommended image capture settings (Q-view imager, Quansys Biosciences).

We verified that cytokine concentrations derived from the Quansys multiplex array were comparable between *Mus* and *Acomys* by testing for parallelism of the mouse standards with *Acomys* serum and tissue lysate. We also evaluated the peptide-level similarity between *Mus* and *Acomys* for each gene represented on the array. Parallelism was examined using standard protocols [92, 93]. Briefly, samples from species and source were randomly pooled to provide a representative cytokine concentration and were run in triplicate at serial dilutions (1:2, 1:6, 1:18, 1:54, and 1:162). To determine parallelism, linear regressions were calculated for samples that had at least 3 dilutions above the lower limit of detection and compared the slopes to the standard curve. For peptide comparisons, the *A. cahirinus* genomic and/or transcribed sequences corresponding to the 16 cytokines of interest were identified by using TBLASTX with inputted *M. musculus* peptide sequence into previously published spiny mouse transcriptomes [40, 94] and an unpublished draft genome. The mRNA sequence was then translated and aligned to peptide sequences for *M. musculus*, *Rattus norvegicus* and *Homo sapiens* using MAFFT [95, 96]. Total similarity and identity was calculated using the Sequence Identity and Similarity (SIAS) tool (http://imed.med.ucm.es/Tools/sias.html).

Individual cytokine concentrations were obtained using image analysis software (Q-view v3.09, Quansys Biosciences). First, the standard curve pixel intensity values were observed and pixel intensity values greater than 60,000 were masked to remove saturated data points. Sample concentrations were calculated from standard curves created by a five parameter logistic regression (5PL) with √y weighting. The average value from each duplicate was then used for subsequent analyses. If the average value was above the lower limit of detection and the pixel intensity co-efficient of variation between duplicates was greater than 15%, the sample was re-assayed on another plate and a new average calculated. Initially, we re-assayed tissues samples below the limit of detection with a greater amount of total protein, but in most cases, additional protein did not equate to quantifiable antigen, suggesting that there was a minimal amount of antigen in those samples. Thus, to maximize use of the plates, we opted to quantify a greater total number of samples and assayed each sample at one dilution. Antigens below the lower limit of detection were recorded as “not present”, and to calculate ratios they were assigned the largest value of the lower limit of detection for that antigen across all plates assayed [97].

### Cox-2 inhibition

Mm-UKY were subjected to a routine ear punch assay and randomly split into two groups: A) 100 mg/kg celecoxib, a potent and specific COX-2 inhibitor or B) vehicle. A Celecoxib capsule was opened and mixed into 0.5% (w/v) methyl cellulose to the appropriate concentration and a 200 µL dose was administered (100 mg active drug / kg body weight) by oral gavage using a 20×30 mm gavage needle tipped with a sugar solution each morning beginning 1 day before injury through 20 days after injury. Ear holes were measured and ear hole area was calculated for every 5 days post injury, as previously described [40]. On D10 and 64 entire ears were harvested from a different set of animals and used for histology and stained with Mason’s Trichrome or Picrosirius red, as previously described [40]. Re-epithelialization was confirmed by the presence of a connected and complete epidermis distal to the amputation plane by examining two tissue sections from the proximal and distal wound sites for each animal at D10. Fibrosis was determined by quantifying the area of collagen deposition in the dermis distal from the amputation plane from two sections from the proximal and distal wound sites using circular polarized light microscopy and the thresholding function in Image J after removing the epidermis, epidermal appendages and tissue artifacts.

### Flow Cytometry

To quantify the number of CD3+ cells present in healing ear tissue, tissue was harvested from a separate group of Mm-UKY and Ac females at D0, 1, 3, 7 and 15 using an 8 mm biopsy punch. Harvested tissue from both ears was combined and a single-cell suspension was created using combination of enzymatic and mechanical digestion, as previously described [15]. Total cells were counted by hemacytometer and incubated with PE-conjugated-anti-CD3 (Clone 17A2, BioLegend, San Diego, CA) at a concentration of 1 µg / 10^6^ cells for 1 hour at room temperature, washed and suspended in cell staining buffer (Cat#420201, BioLegend). Flow cytometry was carried out at the University of Kentucky Flow Cytometry Core using the iCyt Synergy sorter system (Sony Biotechnology Inc., San Jose, CA). Laser calibration and compensation was performed for each experiment using unstained and single fluorescent control samples. Analysis was done using FlowJo (Version 10, FlowJo, LLC, Ashland, OR) to identify CD3-positive lymphocytes by PE fluorescence and forward- and side-scatter. The same gating strategies between species were used (n = 4 or 5 animals per timepoint).

### Immunohistochemistry

To identify the locations of STAT3 responsive cells and CD3+ cells, tissue sections were de-paraffinized, rehydrated, and prepared for examination by light- or fluorescent-microscopy, respectively. For light-microscopy, resident peroxidase was quenched by H_2_O_2_, antigens were exposed by heat-mediated retrieval with sodium citrate buffer, pH=6.0, blocked with serum, incubated with primary antibody (anti-pSTAT3, Cell Signaling Technology Cat#9145, 1:200) overnight at 4°C, incubated with a horseradish peroxidase conjugated secondary (Santa Cruz Biotechnology, Cat#sc-2030, 1:1000) for 1 hour at room temp, treated with 3,3’-Diaminobenzidine (SK-4100, Vector Laboratories, Burlingame, CA) until a visible brown precipitate was observed, counter-stained with hematoxylin, dehydrated and cover-slipped. For fluorescent-microscopy, antigens were exposed by heat mediated retrieval with sodium citrate buffer, pH=6.0, resident avidin and biotin was blocked, sections blocked with serum, incubated with primary antibody (anti-CD3, DAKO, Cat#A0452, 1:500) overnight at 4°C, incubated with a biotin conjugated secondary antibody (Vector Laboratories, Cat#PK-6101, 1:400) for 1 hour at room temp, incubated with streptavidin conjugated AlexaFlour-594 (Molecular Probes, Cat#S11227, 1:5,000), counter-stained with 4’,6-Diamidino-2-Phenylindole, Dihydrochloride (Molecular Probes, Cat#D1306, 1:10,000) and cover-slipped. Images were acquired using a compound epi-fluorescence microscope (IX-51, Olympus Corporation, Tokyo, Japan) equipped with a CCD camera (DP-74, Olympus Corporation) and software (cellSens v1.12, Olympus Corporation).

### Immunoblot

To quantify the STAT3 response to injury, 30 or 40 µg of total protein from tissue lystate was denatured and separated by SDS polyacrylamide gel electrophoresis and transferred to a PVDF membrane (IB401002, Life Technologies). Membranes were cut along the 55 kDa ladder marker and blocked with either 5% BSA or 5% dry skim milk in TBST for 1 hour at room temperature, incubated with primary antibody (pSTAT3, Cell Signaling, Cat#9145, 1:2000; ACTB, Cell Signaling, Cat#4967, 1:5000), washed, incubated with secondary antibody (Santa Cruz Biotechnologies, Cat#sc-2030, 1:10,000), and visualized by chemiluminescence (Cat#RPN2235, GE Healthcare) using a digital CCD camera (UVP LLC, Upland, CA). Total pixel intensity was quantified using regions of interest and normalized to background and uninjured tissue using ImageJ2 [98].

### Statistical Analysis

To compare the baseline serum cytokine concentrations we used a one-way ANOVA and Tukey-Kramer HSD posthoc tests to test for individual group differences. To compare the dynamics of cytokine concentration over time in serum and tissue, a ratio of the injured concentration mean to the uninjured concentration mean was calculated for each cytokine by group (Mm-UKY, Mm-Kenya, Mm-Wild, Ac and Ap) and time point (D1-D20). To normalize cytokine fold-change distributions, data were log transformed. A two-way ANOVA was then used to test for effects of time and group on tissue and serum separately. Pairwise comparisons were tested using the Tukey-Kramer HSD method. In the event that several undetected values existed at an individual timepoint and log transformed data still did not meet normality, we used non-parametric Wilcoxon rank sum tests with Steel Dwass post-hoc tests for pairwise comparisons. Datasets for which non-parametric analyses were performed are indicated in figure legends.

To compare the immunoblot data, pixel intensity was calculated for the bands of interest using an identical sized region of interest with ImageJ [98]. The pixel intensity of pSTAT3 was normalized to ACTB and a two-way ANOVA with time and species was used to compare values and pairwise comparisons were made using the Tukey-Kramer HSD method. To compare the flow cytometry results, we used a two-way ANOVA with time and species on log-transformed data and pairwise comparisons were made using the Tukey-Kramer HSD method. To compare the ear-hole closure rate between control and celecoxib-treated animals we used a repeated-measures ANOVA and cubic regression, as previously published [40]. To compare the ear-hole area and area of tissue positive for Picrosirius, we used a student’s t-test. All statistical tests were done using JMP Pro 14 (SAS Institute Inc., Cary, NC) or Prism 5.0 (GraphPad Software, Inc., San Diego, CA). A *P*-value < 0.05 was used to determine significance for each test. All graphs were created in Prism 5.0 and placed into figures using Illustrator CS5 (Adobe Systems, Inc. San Jose, CA).

## SUPPLEMENTAL INFORMATION

**Supplemental Table 1:**
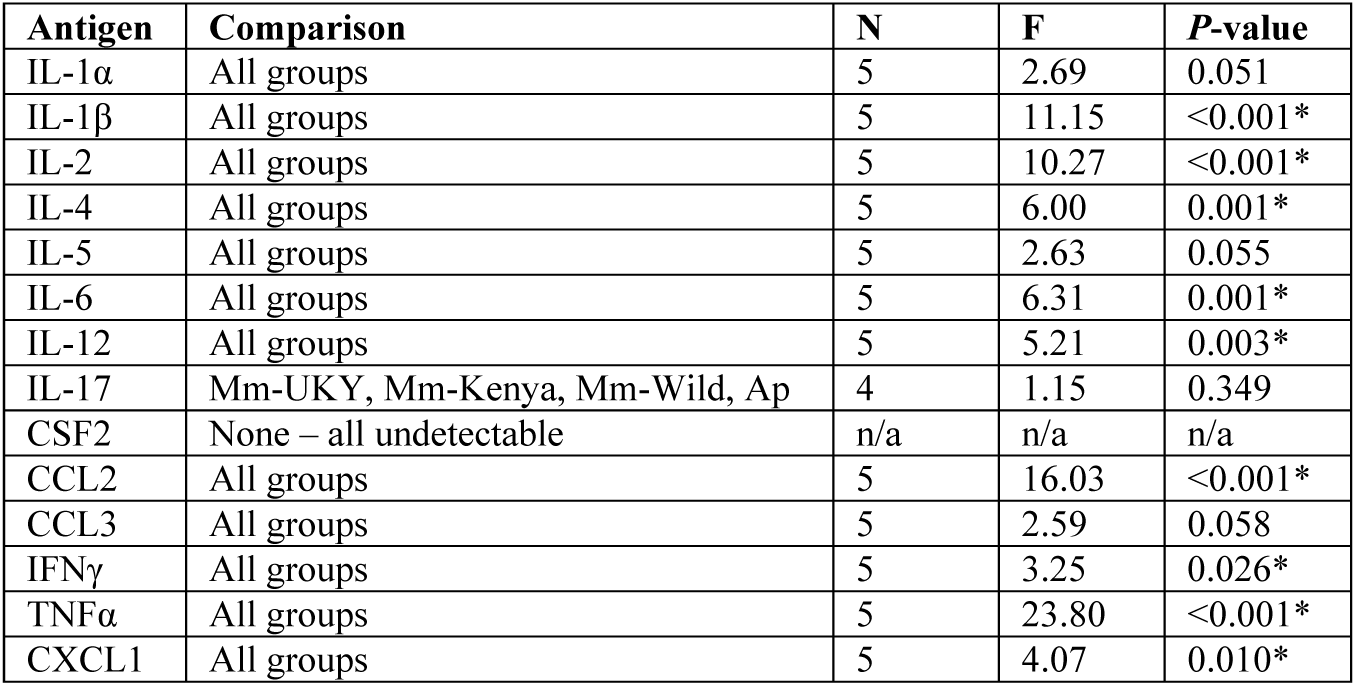
One-way ANOVA analyses of log-transformed uninjured serum data. Groups listed in comparison column represent groups for which data was quantified and could be compared. *P*-values indicate where at least one group is significantly different from another group. Tukey-Kramer HSD post-hoc tests were used for pairwise comparisons are summarized in Figure 1A (See Supplemental File).

| Antigen | Comparison | N | F | *P*-value |
| --- | --- | --- | --- | --- |
| IL-1α | All groups | 5 | 2.69 | 0.051 |
| IL-1β | All groups | 5 | 11.15 | <0.001* |
| IL-2 | All groups | 5 | 10.27 | <0.001* |
| IL-4 | All groups | 5 | 6.00 | 0.001* |
| IL-5 | All groups | 5 | 2.63 | 0.055 |
| IL-6 | All groups | 5 | 6.31 | 0.001* |
| IL-12 | All groups | 5 | 5.21 | 0.003* |
| IL-17 | Mm-UKY, Mm-Kenya, Mm-Wild, Ap | 4 | 1.15 | 0.349 |
| CSF2 | None – all undetectable | n/a | n/a | n/a |
| CCL2 | All groups | 5 | 16.03 | <0.001* |
| CCL3 | All groups | 5 | 2.59 | 0.058 |
| IFNγ | All groups | 5 | 3.25 | 0.026* |
| TNFα | All groups | 5 | 23.80 | <0.001* |
| CXCL1 | All groups | 5 | 4.07 | 0.010* |

**Supplemental Table 2:** Two-way ANOVA analyses of log-transformed serum time series data. Groups listed in comparison column represent groups for which data was quantified throughout the time series. Tukey-Kramer HSD post-hoc tests were used for pairwise comparisons and the results are summarized in Figure 1B (See Supplemental File).

| Antigen | Comparison | N | Group |  | Day |  | Group*Day |  |
| --- | --- | --- | --- | --- | --- | --- | --- | --- |
|  |  |  | F | P-value | F | P-value | F | P-value |
| IL-1α | All groups | 255 | 32.51 | <0.001* | 0.93 | 0.471 | 1.62 | 0.039* |
| IL-1β | Mm-UKY, Mm-Wild, Ap | 157 | 3.02 | 0.052 | 1.06 | 0.391 | 1.08 | 0.386 |
| IL-2 | All groups | 255 | 27.82 | <0.001* | 2.46 | 0.025* | 1.96 | 0.006* |
| IL-4 | All groups | 255 | 34.54 | <0.001* | 1.47 | 0.188 | 1.55 | 0.053 |
| IL-5 | All groups | 255 | 63.22 | <0.001* | 0.94 | 0.469 | 0.82 | 0.706 |
| IL-6 | All groups | 255 | 64.61 | <0.001* | 2.29 | 0.037* | 2.63 | <0.001* |
| IL-12 | All groups | 255 | 33.25 | <0.001* | 2.21 | 0.660 | 0.91 | 0.583 |
| IL-17 | Mm-UKY, Mm-Kenya, Mm-Wild, Ap | 205 | 7.64 | <0.001* | 2.78 | 0.013* | 2.60 | <0.001* |
| CSF2 | None – all undetectable | 0 | n/a | n/a | n/a | n/a | n/a | n/a |
| CCL2 | All groups | 255 | 5.72 | <0.001* | 1.46 | 0.193 | 1.16 | 0.278 |
| CCL3 | All groups | 255 | 38.75 | <0.001* | 0.91 | 0.486 | 1.47 | 0.079 |
| IFNγ | All groups | 255 | 37.81 | <0.001* | 1.22 | 0.296 | 1.77 | 0.018* |
| TNFα | All groups | 255 | 31.88 | <0.001* | 0.85 | 0.535 | 0.91 | 0.583 |
| CXCL1 | Mm-UKY, Mm-Kenya, Mm-Wild, Ap | 205 | 69.43 | <0.001* | 4.33 | <0.001* | 1.47 | 0.104 |

**Supplemental Table 3:** Two-way ANOVA analyses of log-transformed tissue lysate time series data. Groups listed in comparison column represent groups for which data was quantified throughout the time series. *P*-values indicate where at least one group is significantly different from another group. Tukey-Kramer HSD post-hoc tests were used for pairwise comparisons and the results are summarized in Figure 2 (See Supplemental File).

| Antigen | Comparison | N | Group |  | Day |  | Group*Day |  |
| --- | --- | --- | --- | --- | --- | --- | --- | --- |
|  |  |  | F | *P*-value | F | *P*-value | F | *P*-value |
| IL-1α | None – all above limit of detection | 0 | n/a | n/a | n/a | n/a | n/a | n/a |
| IL-1β | All groups | 261 | 77.88 | <0.001* | 39.47 | <0.001* | 5.33 | <0.001* |
| IL-2 | All groups | 261 | 190.93 | <0.001* | 6.68 | <0.001* | 13.59 | <0.001* |
| IL-4 | All groups | 261 | 29.09 | <0.001* | 11.24 | <0.001* | 6.76 | <0.001* |
| IL-5 | Mm-UKY, Mm-Kenya, Mm-Wild, Ap | 211 | 22.73 | <0.001* | 10.03 | <0.001* | 10.70 | <0.001* |
| IL-6 | All groups | 261 | 76.89 | <0.001* | 104.11 | <0.001* | 15.06 | <0.001* |
| IL-12 | All groups | 261 | 123.20 | <0.001* | 10.42 | <0.001* | 18.28 | <0.001* |
| IL-17 | Mm-UKY, Mm-Kenya, Ap | 177 | 598.63 | <0.001* | 4.42 | <0.001* | 17.55 | <0.001* |
| CSF2 | Mm-UKY, Mm-Kenya, Mm-Wild, Ap | 211 | 104.44 | <0.001* | 21.36 | <0.001* | 8.51 | <0.001* |
| CCL2 | Mm-UKY, Mm-Kenya, Mm-Wild, Ap | 211 | 72.96 | <0.001* | 38.71 | <0.001* | 5.23 | <0.001* |
| CCL3 | All groups | 261 | 8.30 | <0.001* | 160.11 | <0.001* | 6.24 | <0.001* |
| IFNγ | Mm-Kenya, Mm-Wild, Ap | 161 | 40.47 | <0.001* | 2.42 | 0.030* | 3.00 | <0.001* |
| TNFα | All groups | 261 | 149.87 | <0.001* | 81.32 | <0.001* | 11.84 | <0.001* |
| CXCL1 | Mm-UKY, Mm-Kenya, Mm-Wild, Ap | 211 | 137.76 | <0.001* | 201.75 | <0.001* | 26.64 | <0.001* |

**Supplemental Table 4:**
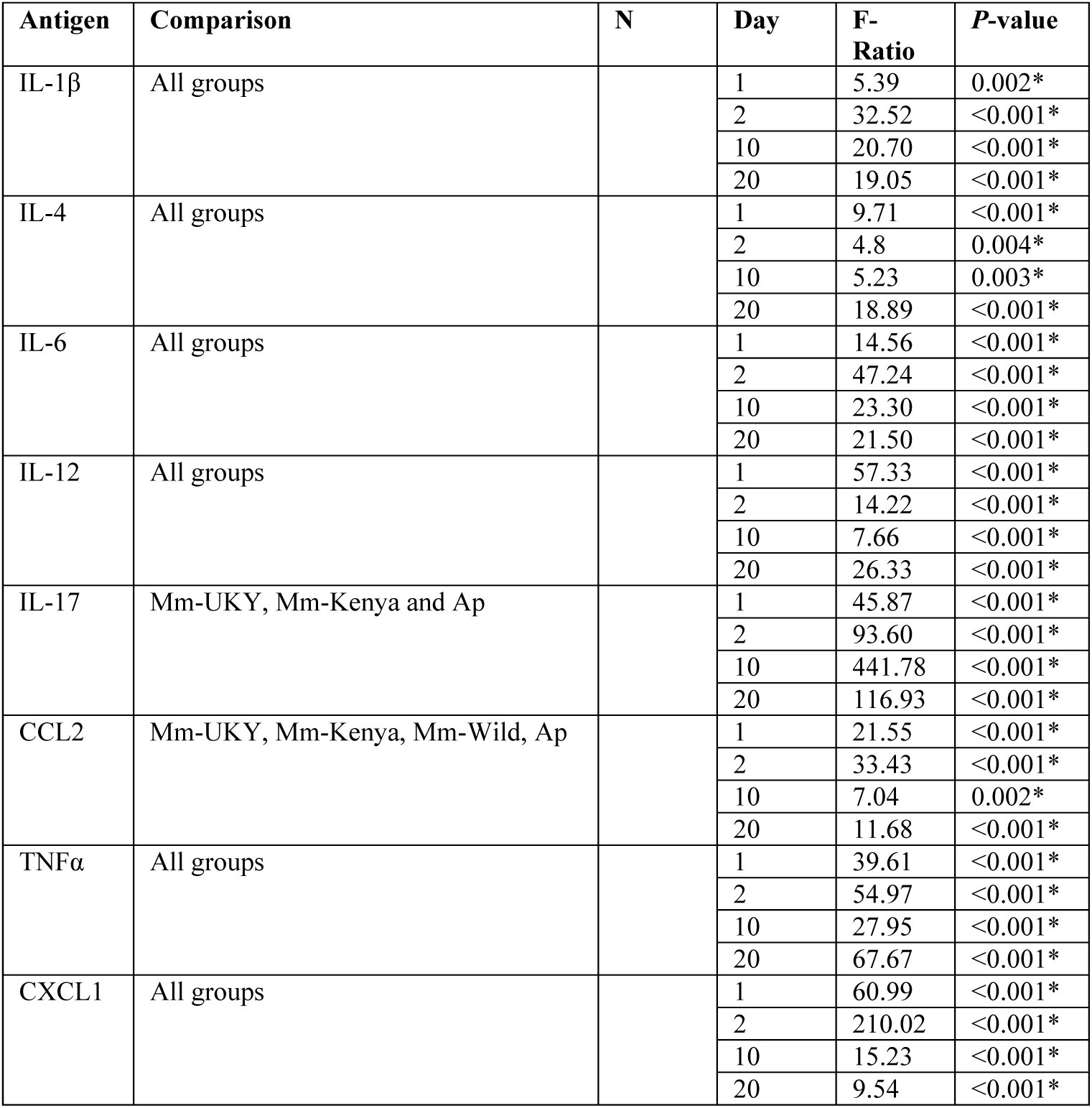
One-way ANOVA analysis of tissue cytokine data to test for an effect of group at D1, 2, 10 and 20. *P*-values indicate where at least one group is significantly different from another group. Tukey-Kramer HSD post-hoc tests were used for pairwise comparisons and are summarized in Figure 3 (See Supplemental File).

| Antigen | Comparison | N | Day | F-Ratio | *P*-value |
| --- | --- | --- | --- | --- | --- |
| IL-1β | All groups | | 1 | 5.39 | 0.002* |
|  |  |  | 2 | 32.52 | <0.001* |
|  |  |  | 10 | 20.70 | <0.001* |
|  |  |  | 20 | 19.05 | <0.001* |
| IL-4 | All groups |  | 1 | 9.71 | <0.001* |
|  |  |  | 2 | 4.8 | 0.004* |
|  |  |  | 10 | 5.23 | 0.003* |
|  |  |  | 20 | 18.89 | <0.001* |
| IL-6 | All groups |  | 1 | 14.56 | <0.001* |
|  |  |  | 2 | 47.24 | <0.001* |
|  |  |  | 10 | 23.30 | <0.001* |
|  |  |  | 20 | 21.50 | <0.001* |
| IL-12 | All groups |  | 1 | 57.33 | <0.001* |
|  |  |  | 2 | 14.22 | <0.001* |
|  |  |  | 10 | 7.66 | <0.001* |
|  |  |  | 20 | 26.33 | <0.001* |
| IL-17 | Mm-UKY, Mm-Kenya and Ap |  | 1 | 45.87 | <0.001* |
|  |  |  | 2 | 93.60 | <0.001* |
|  |  |  | 10 | 441.78 | <0.001* |
|  |  |  | 20 | 116.93 | <0.001* |
| CCL2 | Mm-UKY, Mm-Kenya, Mm-Wild, Ap |  | 1 | 21.55 | <0.001* |
|  |  |  | 2 | 33.43 | <0.001* |
|  |  |  | 10 | 7.04 | 0.002* |
|  |  |  | 20 | 11.68 | <0.001* |
| TNFα | All groups | | 1 | 39.61 | <0.001* |
|  |  |  | 2 | 54.97 | <0.001* |
|  |  |  | 10 | 27.95 | <0.001* |
|  |  |  | 20 | 67.67 | <0.001* |
| CXCL1 | All groups |  | 1 | 60.99 | <0.001* |
|  |  |  | 2 | 210.02 | <0.001* |
|  |  |  | 10 | 15.23 | <0.001* |
|  |  |  | 20 | 9.54 | <0.001* |

**Supplemental Table 5:**
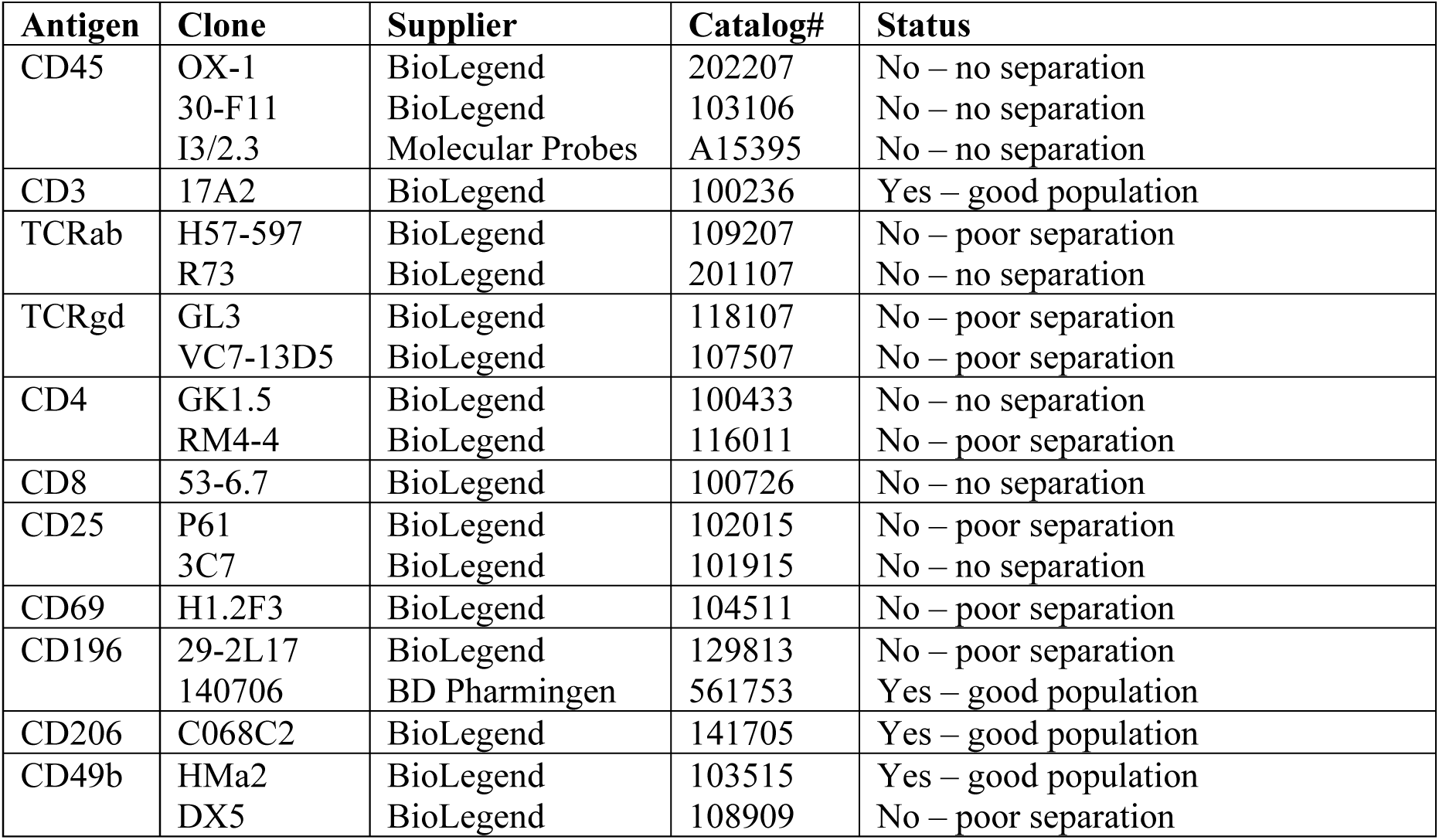
List of T cell phenotyping antibodies tried using flow cytometry in *Acomys cahirinus*.

**Supplemental Figure 1.**
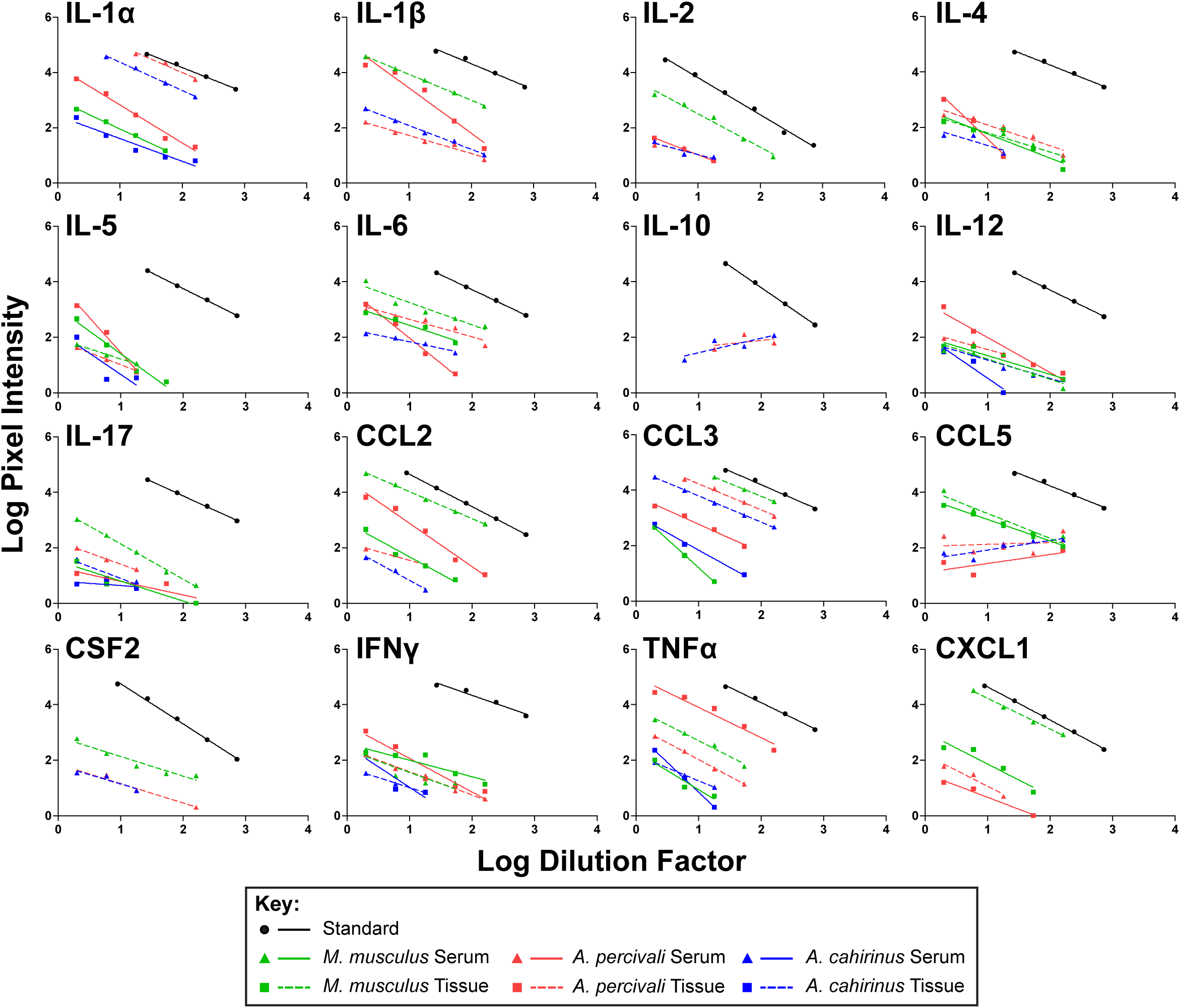
Comparison of parallelism for cytokine assays demonstrates ability to quantify cytokines across three species. Serial dilution of serum (triangles, solid lines) and tissue lystate (squares, dashed lines) shows similar negative slopes for M. musculus (green), A. percivali (red) and A. cahirinus (blue) compared to the standard (black, circles, solid line). Data represent the mean for duplicates and lines are linear regressions calculated from all data that was above the lower limit of detection.

**Supplemental Figure 1a.**
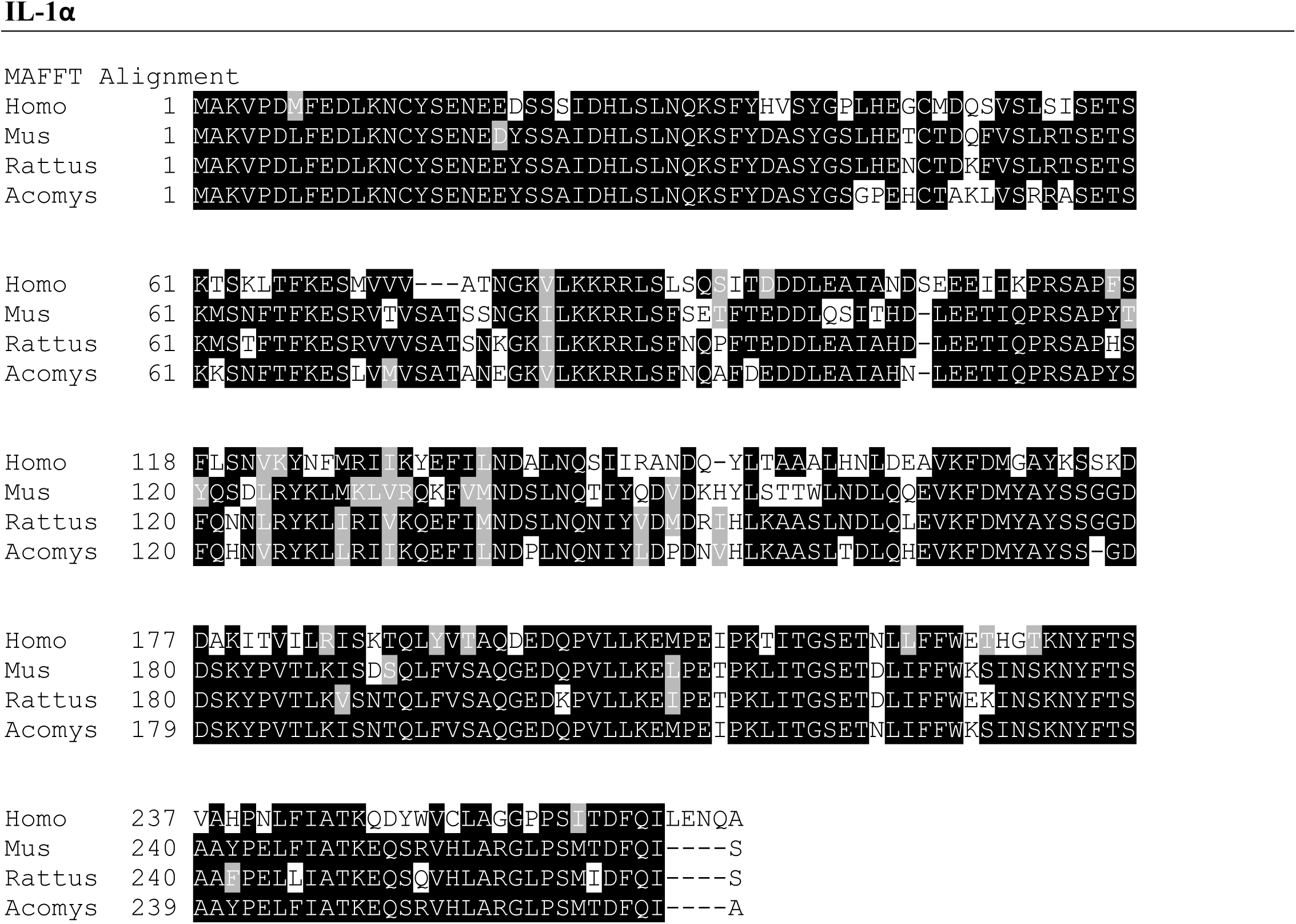

**Supplemental Figure 1b.**
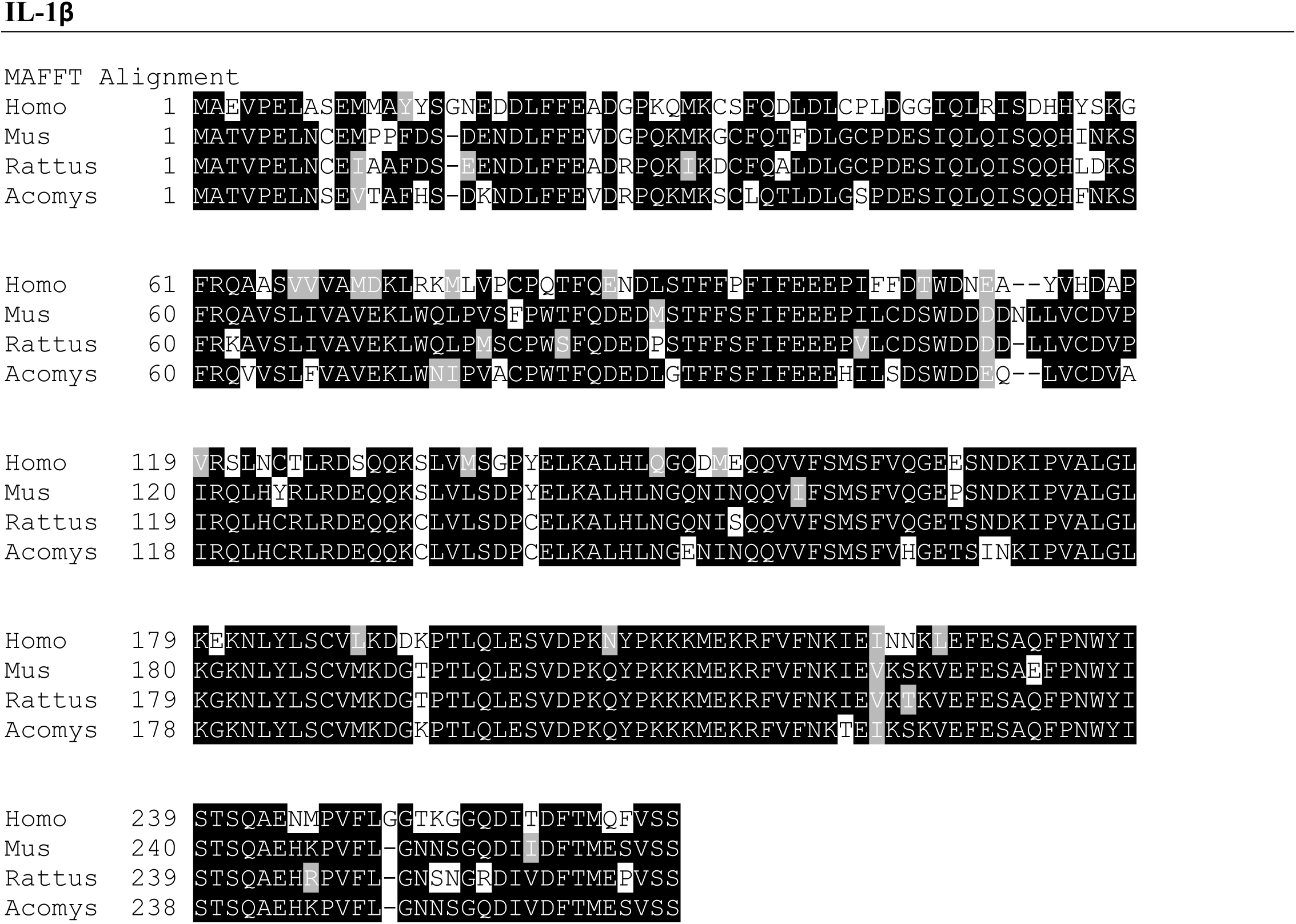

**Supplemental Figure 1c.**
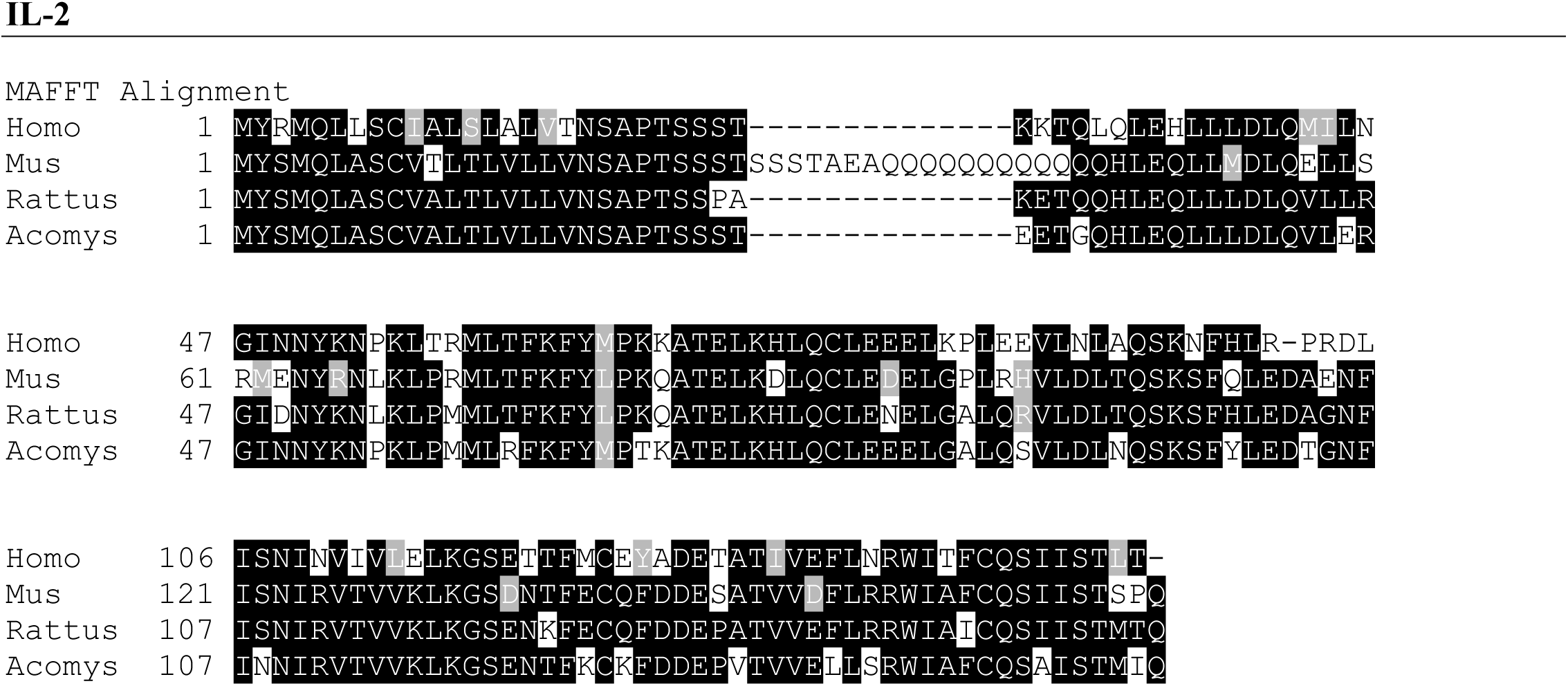

**Supplemental Figure 1d.**
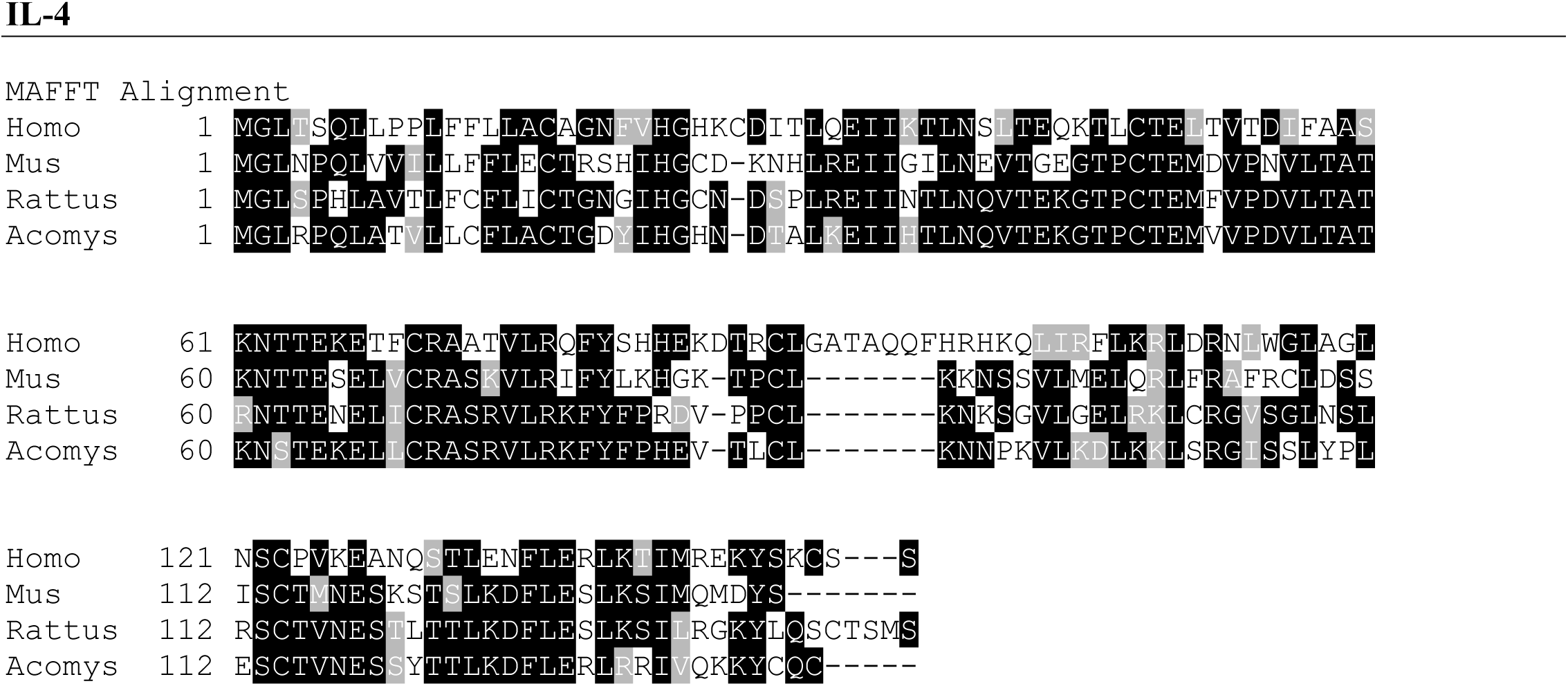

**Supplemental Figure 1e.**
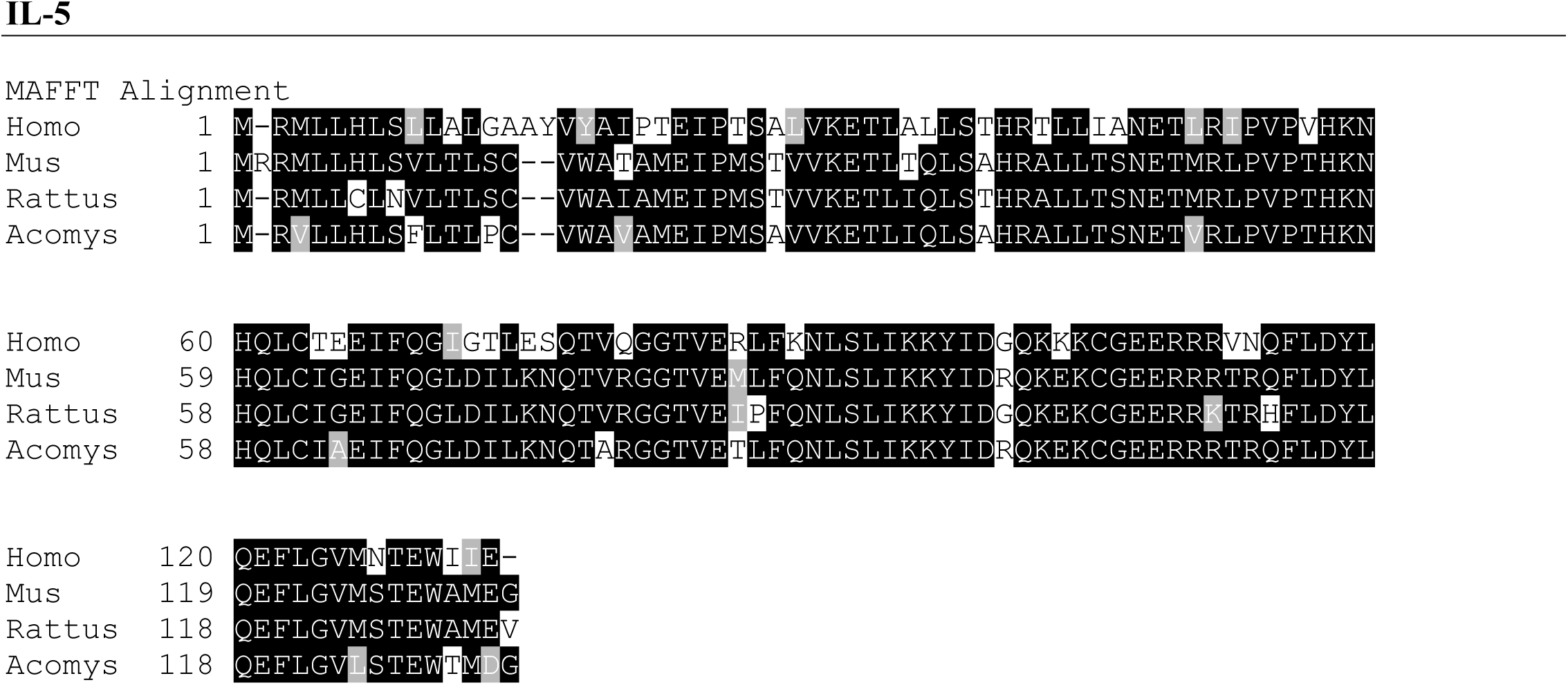

**Supplemental Figure 1f.**
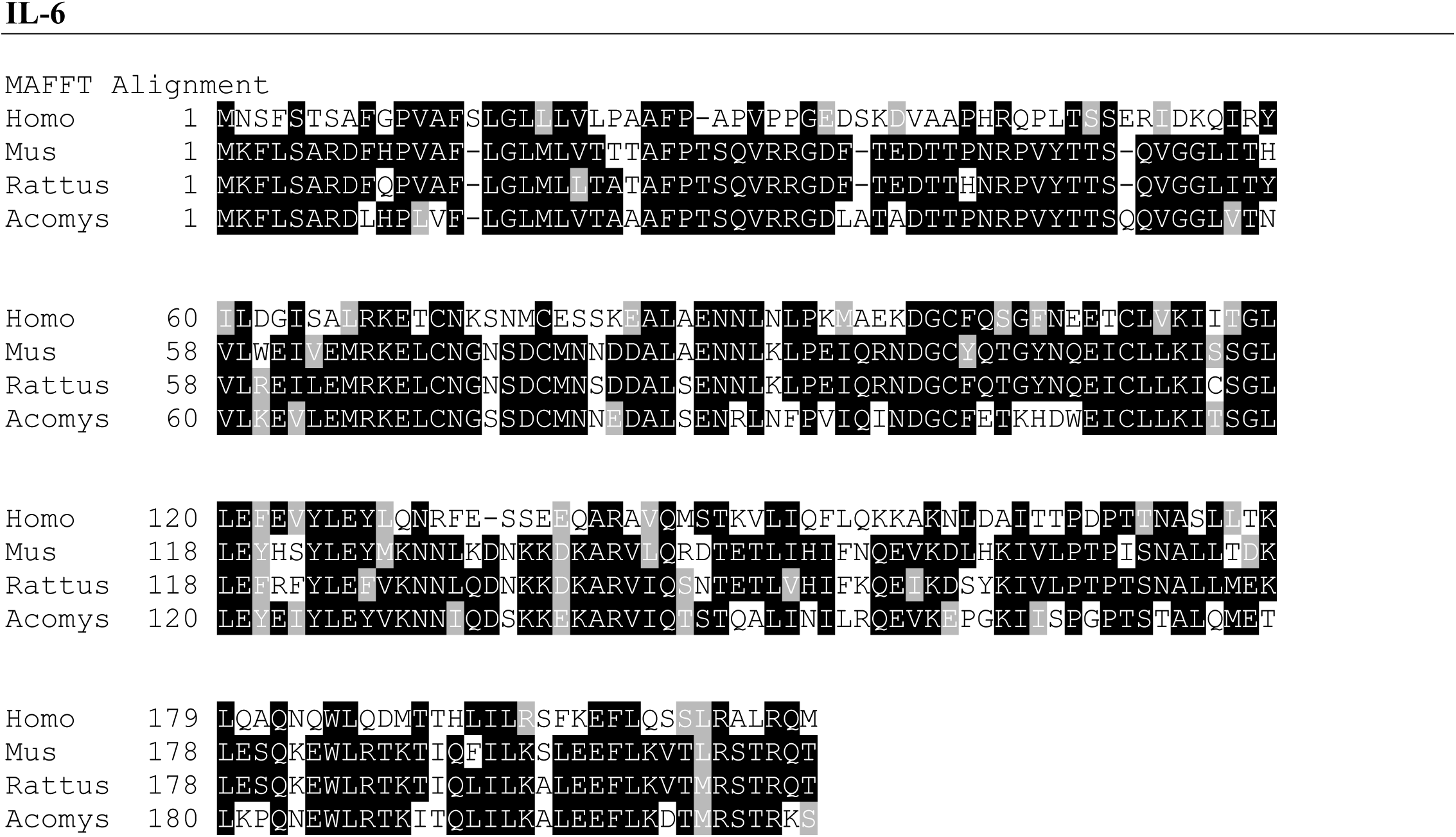

**Supplemental Figure 1g.**
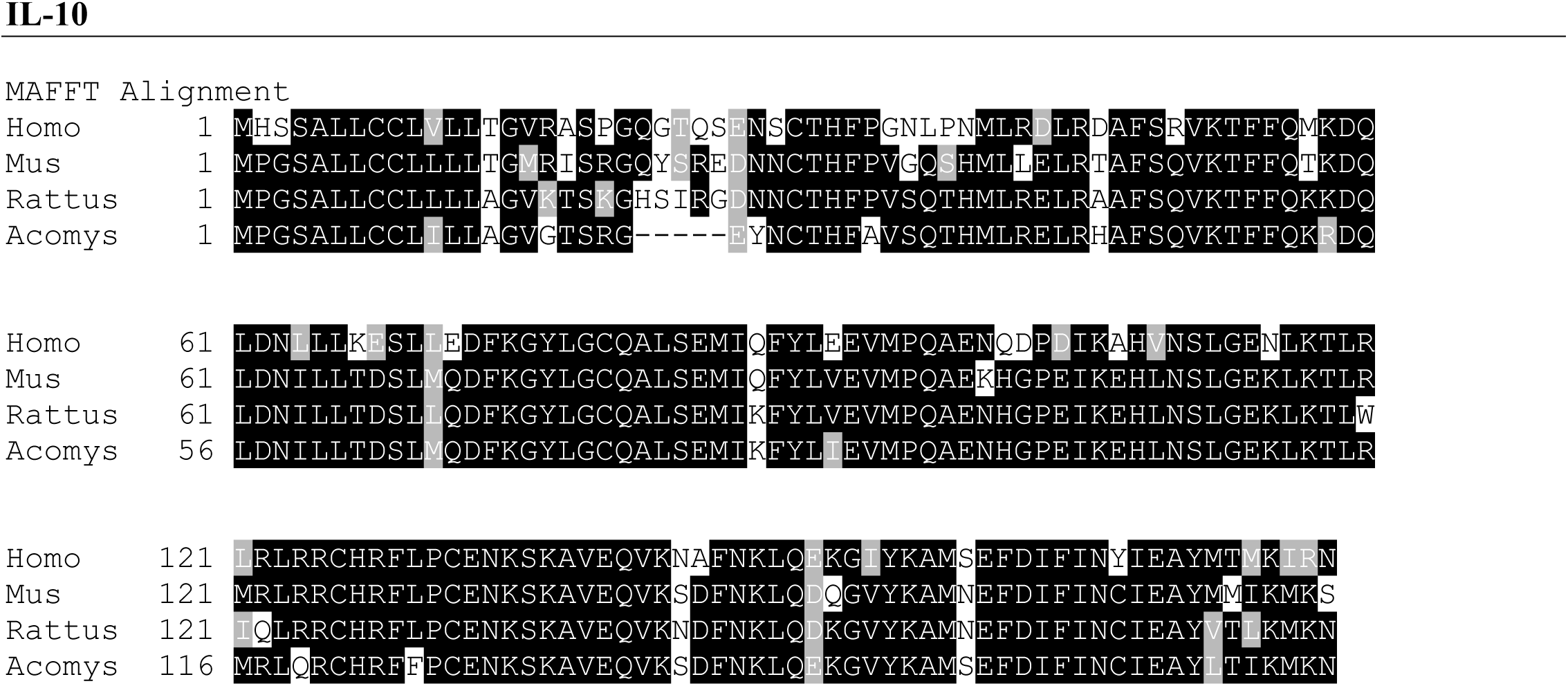

**Supplemental Figure 1h.**
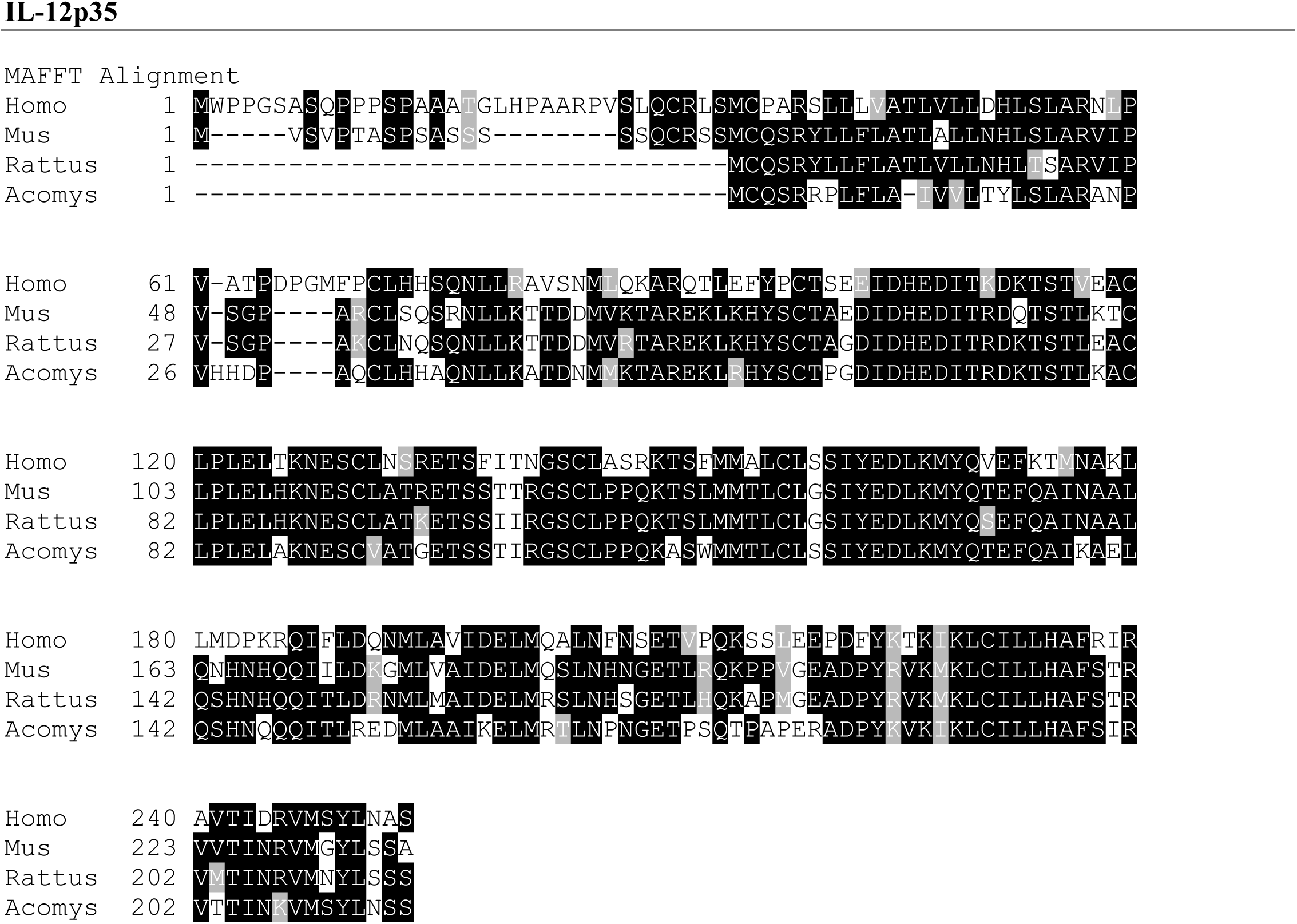

**Supplemental Figure 1i.**
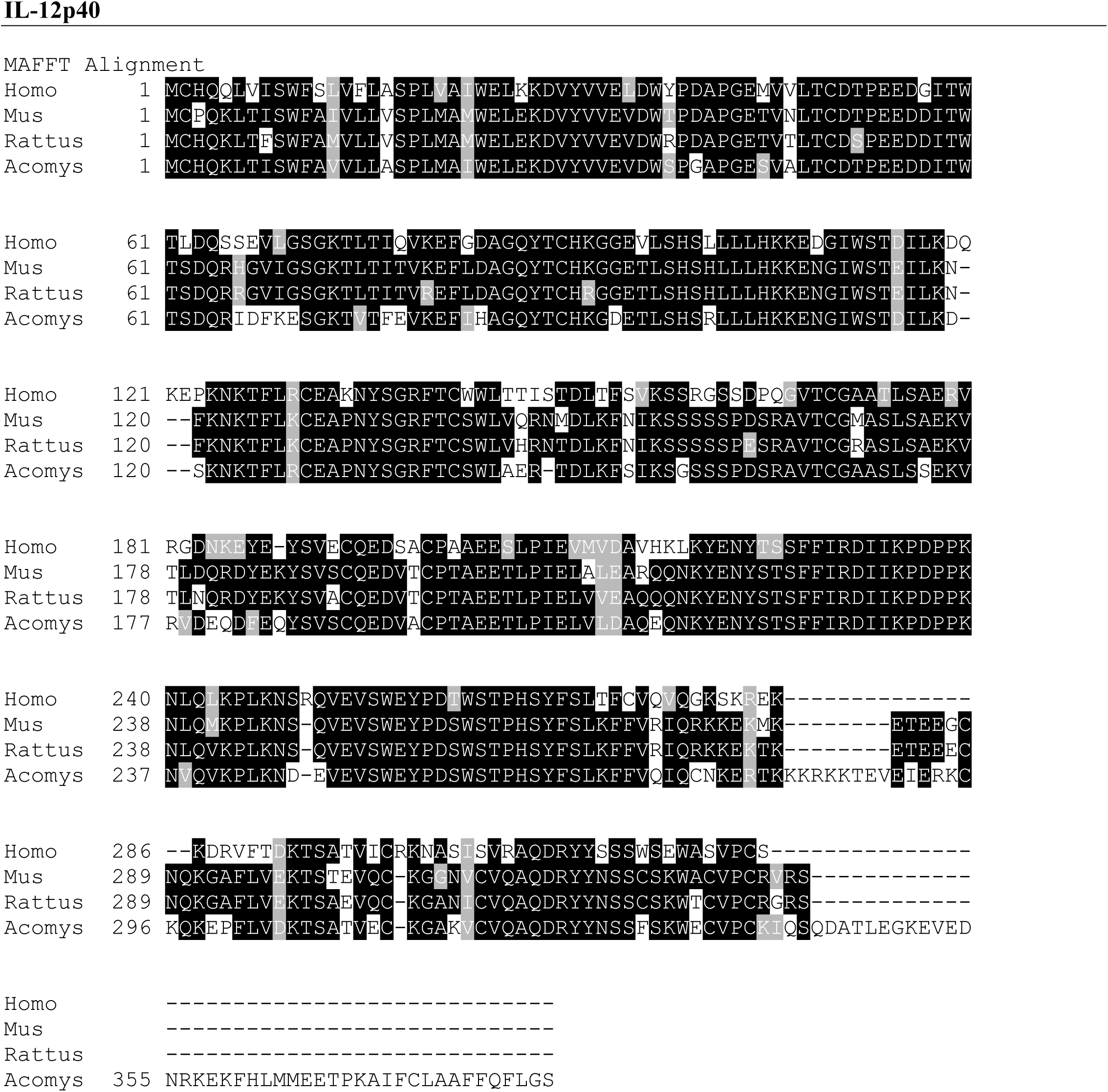

**Supplemental Figure 1j.**
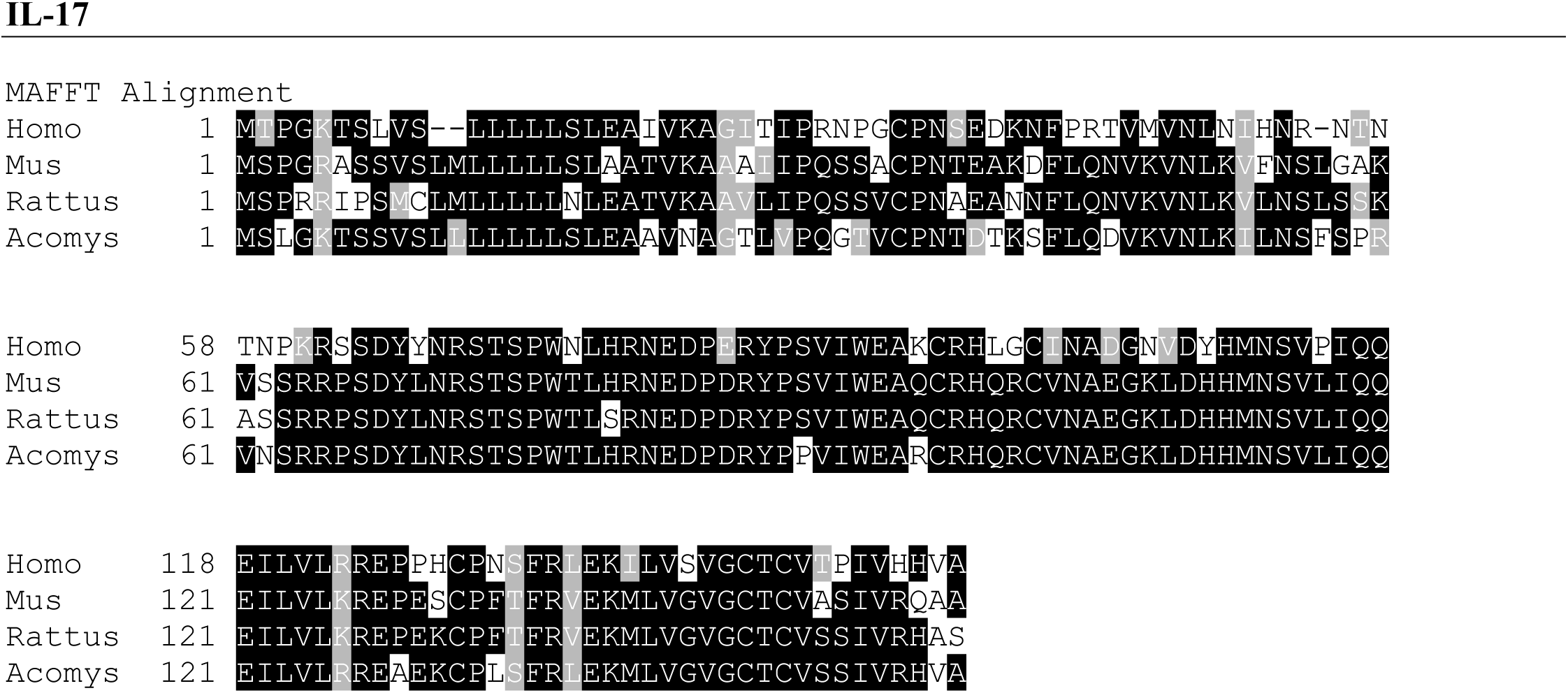

**Supplemental Figure 1k.**
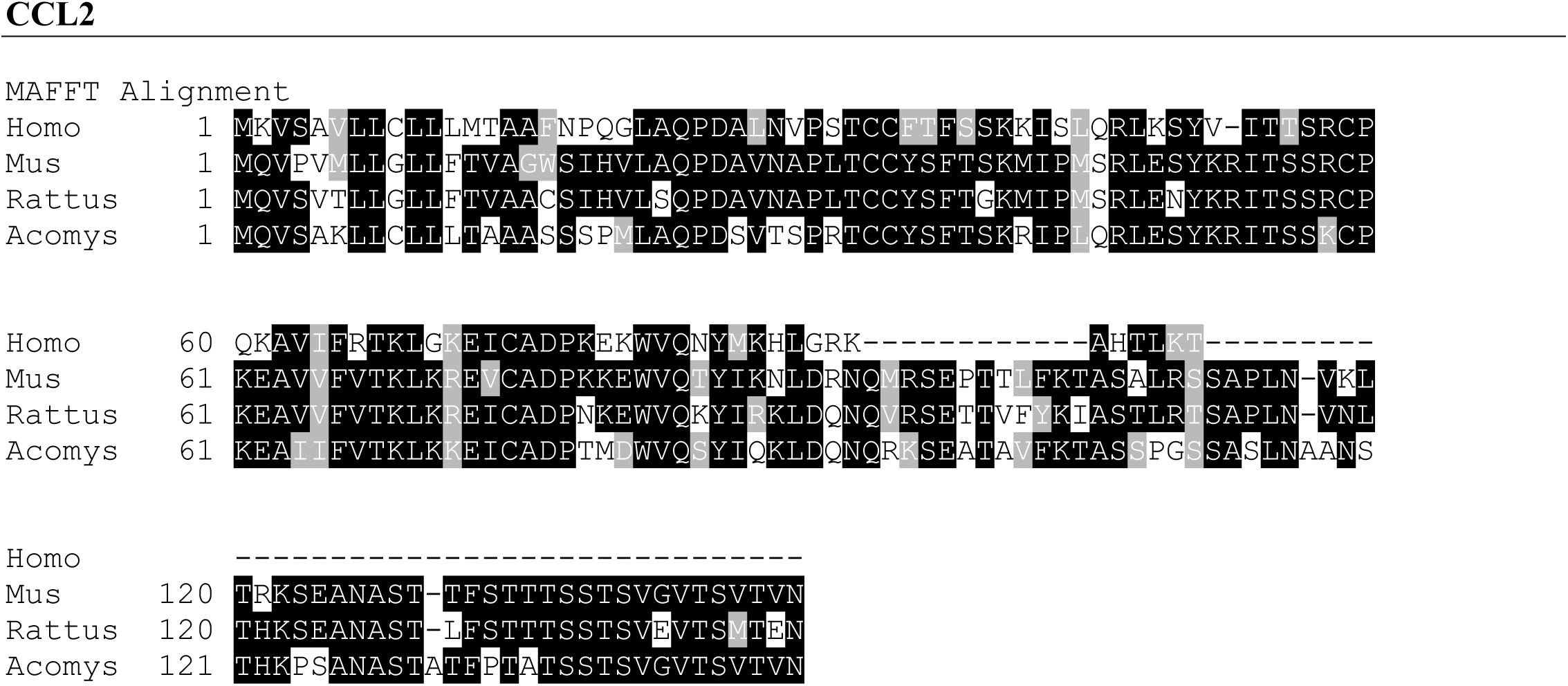

**Supplemental Figure 1l.**
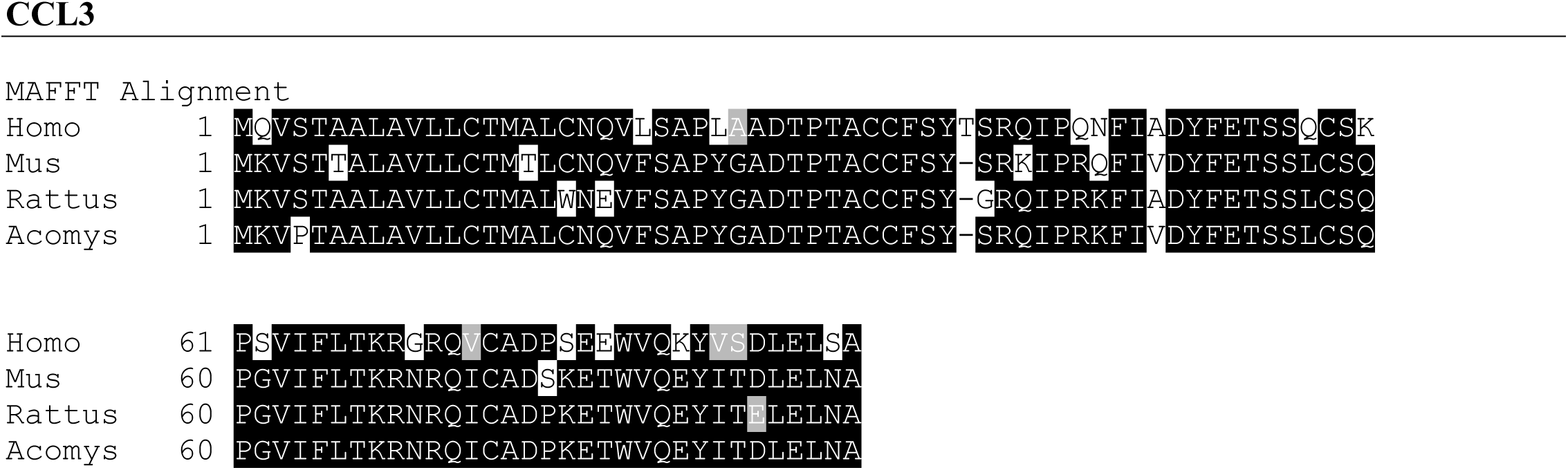

**Supplemental Figure 1m.**
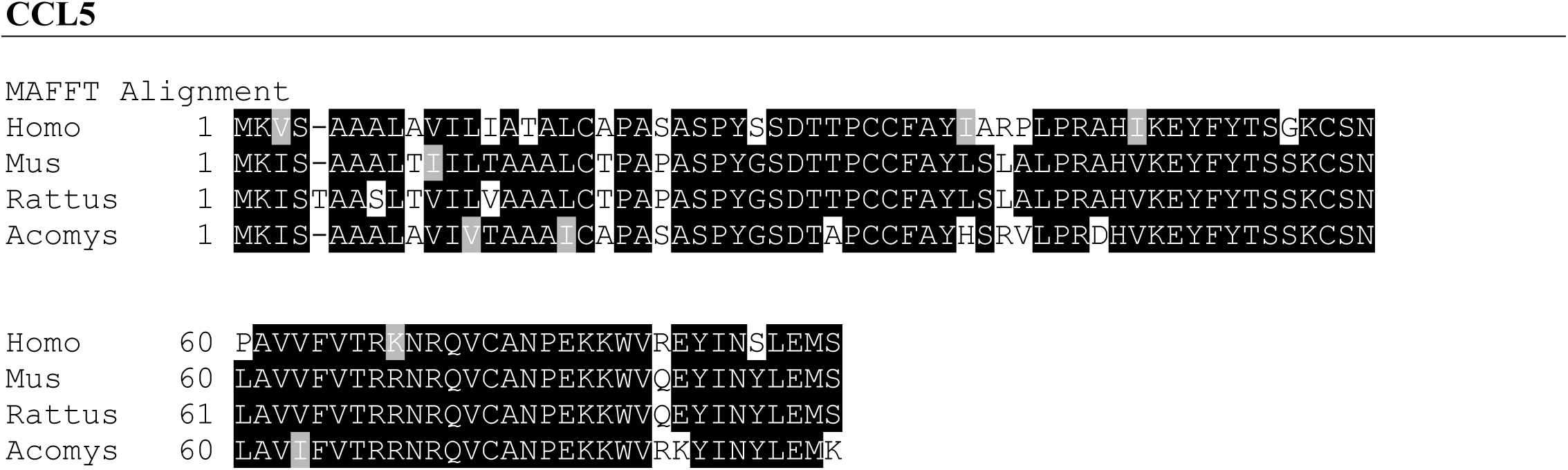

**Supplemental Figure 1n.**
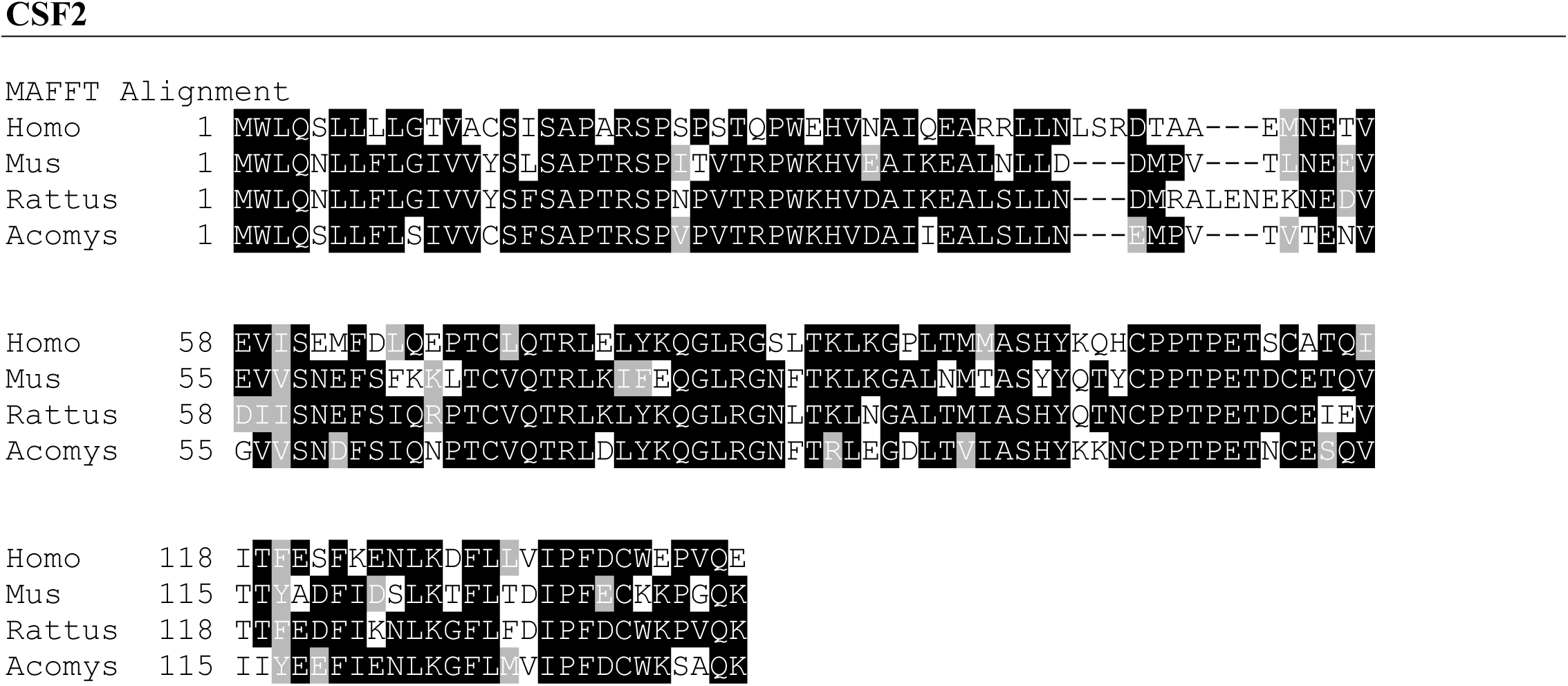

**Supplemental Figure 1o.**
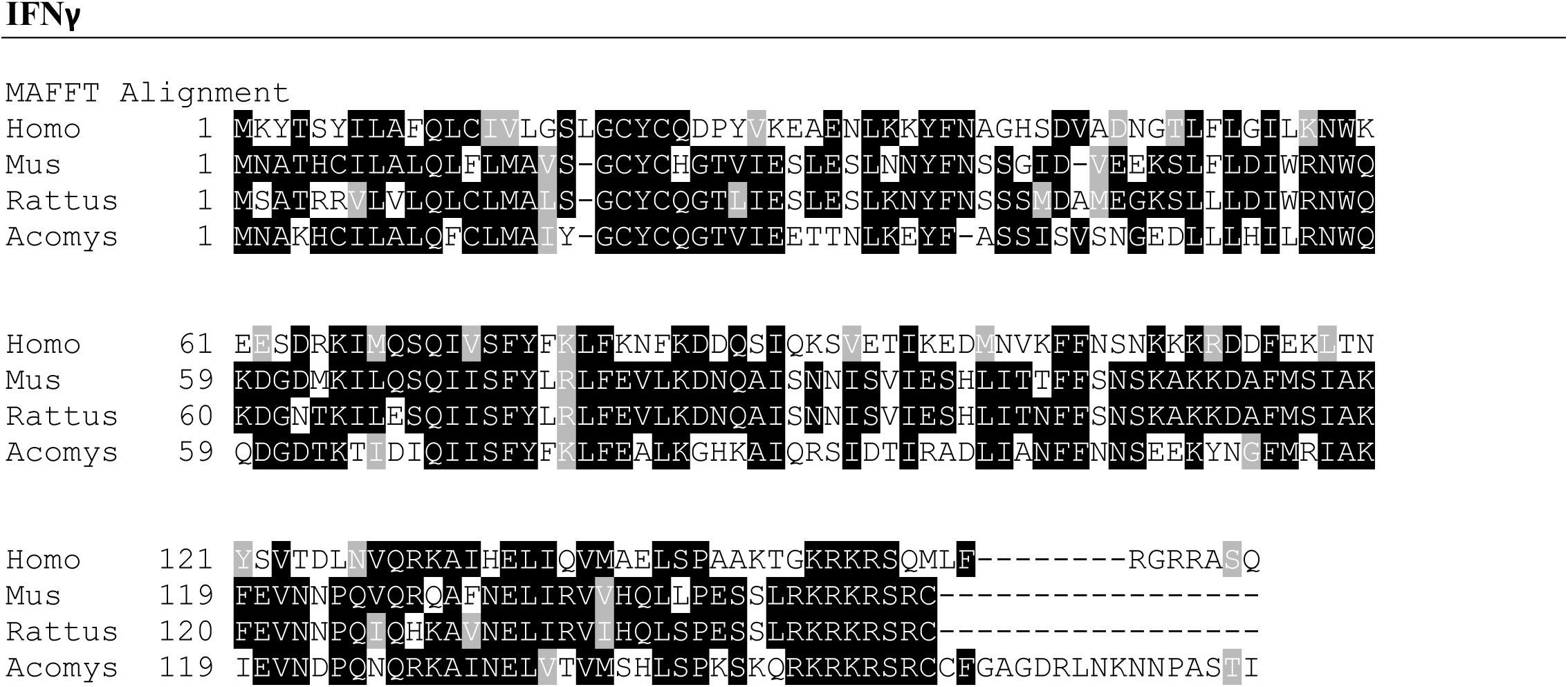

**Supplemental Figure 1p.**
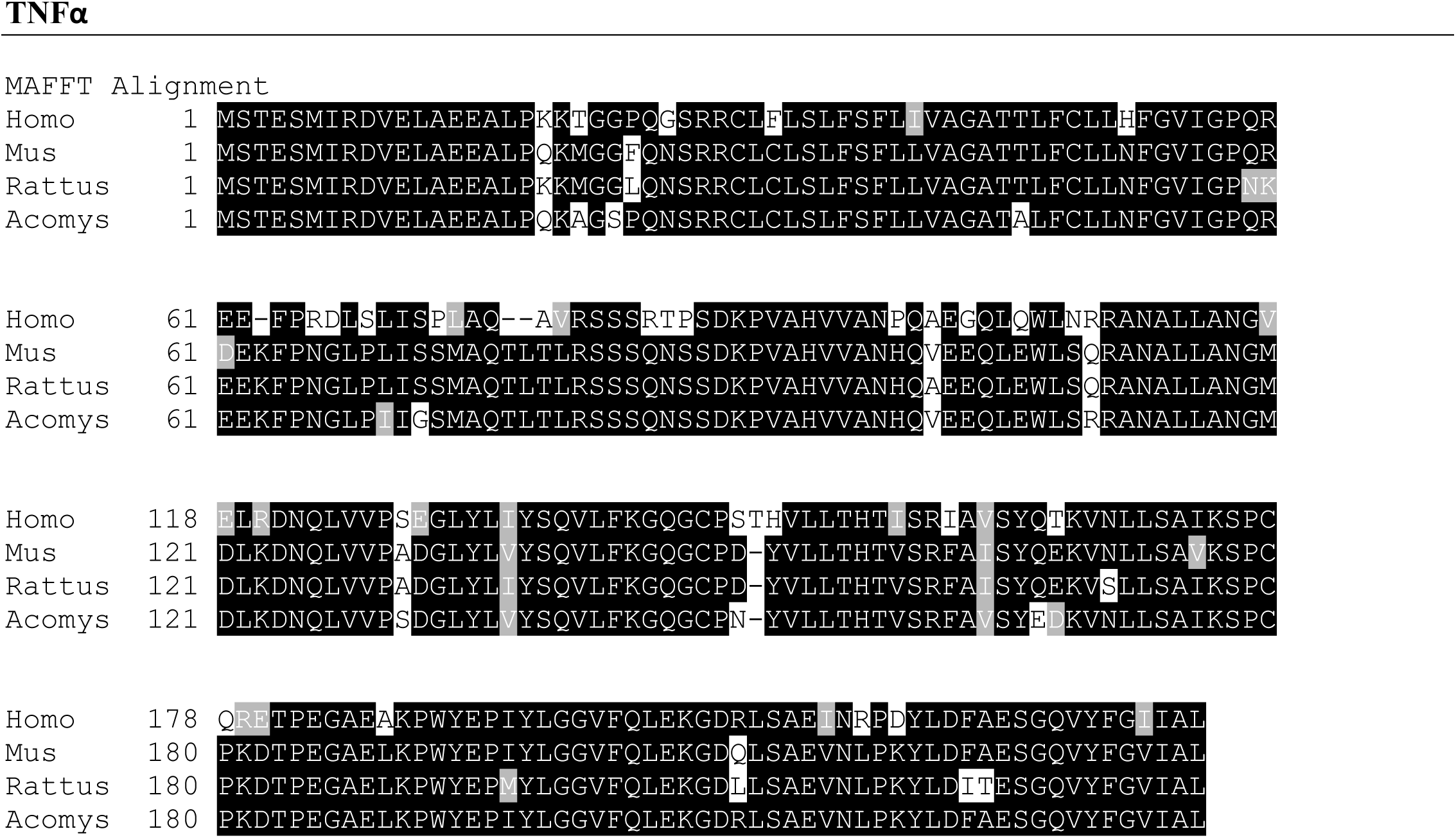

**Supplemental Figure 1q.**
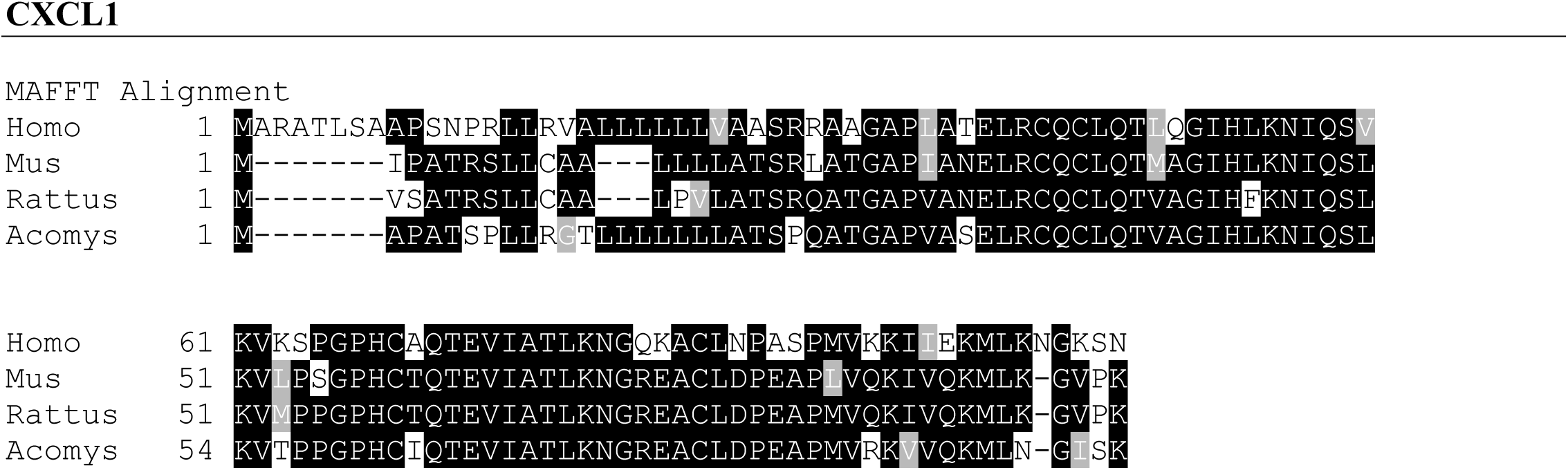

**Supplemental Figure 2 Comparisons of amino acid sequence alignments for antigens in study**. Predicted amino acid sequence was translated from mRNA (*A. cahirinus*) or obtained from NCBI (*H. sapiens*, *M. musculus*, *R. rattus*) and aligned together using MAFFT. Figures were created using BOXSHADE where solid black shading is equivalent, gray shading is similar and no shading is dissimilar.

## Supplemental Files

Supplemental File 1 contains summary tables of post-hoc multiple comparison tests used in Figure 1B, Figure 2 and Figure 3.

